# Mitochondrial phosphate carrier-dependence of mitochondrial calcium chelation and respiration in skeletal muscle

**DOI:** 10.64898/2026.08.03.742530

**Authors:** Cesar Vásquez-Trincado, Valentina Debattisti, Arijita Gosh, Amanda H. Collins, Ji In Han, Carmen Bekeová, Emanuele Loro, Tejvir Khurana, György Hajnóczky, Erin L. Seifert

## Abstract

The past decade has seen remarkable progress in the molecular resolution of the influx and efflux components of mitochondrial Ca^2+^ handling, but little progress in matrix Ca^2+^ chelation that is central to mitochondrial calcium signaling. Among possible chelators, inorganic phosphate (Pi) is dynamic, and its uptake can lessen during Ca^2+^ uptake the rundown of the membrane potential, the primary driving force for Ca^2+^ uptake. Thus, PiC, the major mitochondrial Pi transporter, is well-positioned to regulate mitochondrial Ca^2+^ handling. To test this, we depleted (KD) PiC in murine skeletal muscle. We show that PiC depletion enhances the matrix free Ca^2+^ rise across a range of Ca^2+^ uptake activities, and surprisingly, causes elevated mitochondrial Ca^2+^ uptake, which seems to arise from an increased abundance of the mitochondrial Ca^2+^ uniporter. With protein levels of non-mitochondrial Ca^2+^ handling proteins unaltered, the greater mitochondrial Ca^2+^ uptake may drive a suppressed cytoplasmic Ca^2+^ response to tetanic stimulation, via lesser Ca^2+^-mediated positive feedback on Ca^2+^ release channels, contributing to an exercise deficit in KD mice. Alternatively, less buffering of matrix Ca^2+^ might be beneficial. We test these possibilities by lowering MCU, the uniporter’s pore forming subunit, in skeletal muscle of adult PiC KD mice, and find a worsened exercise deficit. This study establishes the requirement for PiC to maintain a bound fraction of Ca^2+^ in the matrix, and reveals a fitness benefit for elevated [Ca^2+^]m in striated muscle depleted of PiC.

## INTRODUCTION

The role of mitochondria in cellular Ca^2+^ homeostasis has been of interest since it was appreciated that the cytoplasmic Ca^2+^ concentration ([Ca^2+^]_c_) can rise high enough to overcome the low affinity of the mitochondrial Ca^2+^ uniporter (mtCU) in the inner mitochondrial membrane (IMM) (1–5). Since then, several studies have shown a correlation between mitochondrial matrix Ca^2+^ and different aspects of cell function (e.g.,(6–13)). Yet, the mechanisms that underlie these associations are not always evident, or straightforward to test. Understanding how mitochondria handle Ca^2+^ can help to better explain how mitochondrial Ca^2+^ handling is integrated into the broader functions of the cell.

Mitochondrial Ca^2+^ handling has three components - influx, efflux and chelation - and the past decade has seen remarkable progress on the first two. We now know that uptake across the inner mitochondrial membrane (IMM) occurs via a hetero-oligomeric complex (mtCU) formed by a tetrameric pore forming subunit, MCU (14, 15), a scaffold, EMRE, required for function (16), and regulatory subunits (MICUs: MICU1/2/3) that exist as heterodimers and possibly homodimers (17–21), always with MICU1 bound to EMRE and MCU (22–25). An MCU paralog, MCUb, can also be present and produces a dominant negative phenotype (26). On the efflux side, Na^+^-dependent and independent exchangers exist in the IMM (27, 28). NCLX was identified as a mitochondrial Na^+^-Ca^2+^ exchanger (29), and TMBIM5 was recently proposed as a H^+^-Ca^2+^ exchanger (30, 31), but needs further validation.

Concerning Ca^2+^ chelation in the mitochondrial matrix, the vast majority of matrix Ca^2+^ appears to exist in the bound form (estimated bound-to-free of 150,000 in rat brain mitochondria (32) and 800 in rat heart mitochondria (33)), pointing to the importance of enabling mechanisms. Several species, including cardiolipin (CL), ATP and inorganic phosphate (Pi) have relevance. While CL and ATP are fairly stable components of Ca^2+^ binding, chelation by Pi is complex and dynamic in capacity. In vitro, Ca^2+^ and Pi can aggregate and precipitate under permissive conditions, and it was hypothesized in the 1960s that Ca^2+^-Pi aggregates would form in the mitochondrial matrix. Early efforts to detect Ca^2+^-Pi aggregates relied on chemical fixation of samples (34). Recent attempts to visualize Ca^2+^-Pi in cells and isolated mitochondria used vitrification along with FCCP-dependence and measurement of elemental composition, providing compelling evidence that Ca^2+^-Pi aggregates can form in the matrix (35–37). Pi uptake is expected to be largely mediated by a high capacity mitochondrial Pi carrier, PiC (SLC25A3), in the IMM (38–41). Measurements in isolated mitochondria of Ca^2+^ clearance and of concentration of free Ca^2+^ in the matrix ([Ca^2+^]m) with different [Pi] have provided functional evidence that Pi chelates matrix Ca^2+^ (32, 33). Beyond Ca^2+^ chelation, Pi uptake that accompanies Ca^2+^ uptake can mitigate the rundown of the membrane potential, the primary driving force for Ca^2+^ uptake (32, 33, 42). Thus Pi uptake and matrix Pi are positioned to be important regulatory nodes for mitochondrial Ca^2+^handling. Yet, all studies in the molecular era have focused on Ca^2+^ influx and efflux. Here we attempt to fill this gap using a murine model in which PiC is targeted in the skeletal muscle (SM). SM has a high capacity for mitochondrial Ca^2+^ uptake (43, 44) and expression of mtCU components (44, 45).

## RESULTS

### New model of skeletal muscle-specific PiC depletion

Mice harboring floxed PiC alleles (PiC^fl/fl^) (41) were crossed with mice expressing the double fusion protein comprising MerCreMer and the Human Skeletal Actin promoter (HSA-MCM) (46) to generate mice with SM-specific Tamoxifen (Tam)-inducible PiC knockdown (KD). HSA-MCM-PiC^fl/fl^ (KD) and PiC^fl/fl^ (Ctrl) mice were treated with Tam for 4 days starting at 9 weeks of age. Unexpectedly, PiC KD was found to be constitutive, though still confined to SM (see **SFig1A**). Further details of the mouse model can be found in Materials and Methods.

Immunoblot evaluation of PiC protein expression revealed that PiC was depleted by 82-85% in *Soleus* and *Flexor digitorum brevis*, and 90-97% *Extensor digitorum longus* and *Quadriceps* (Quad) (Fig 1A). PiC was depleted in SM and not in heart, brain, liver, or kidney (**SFig 1A**).

**Figure 1:**
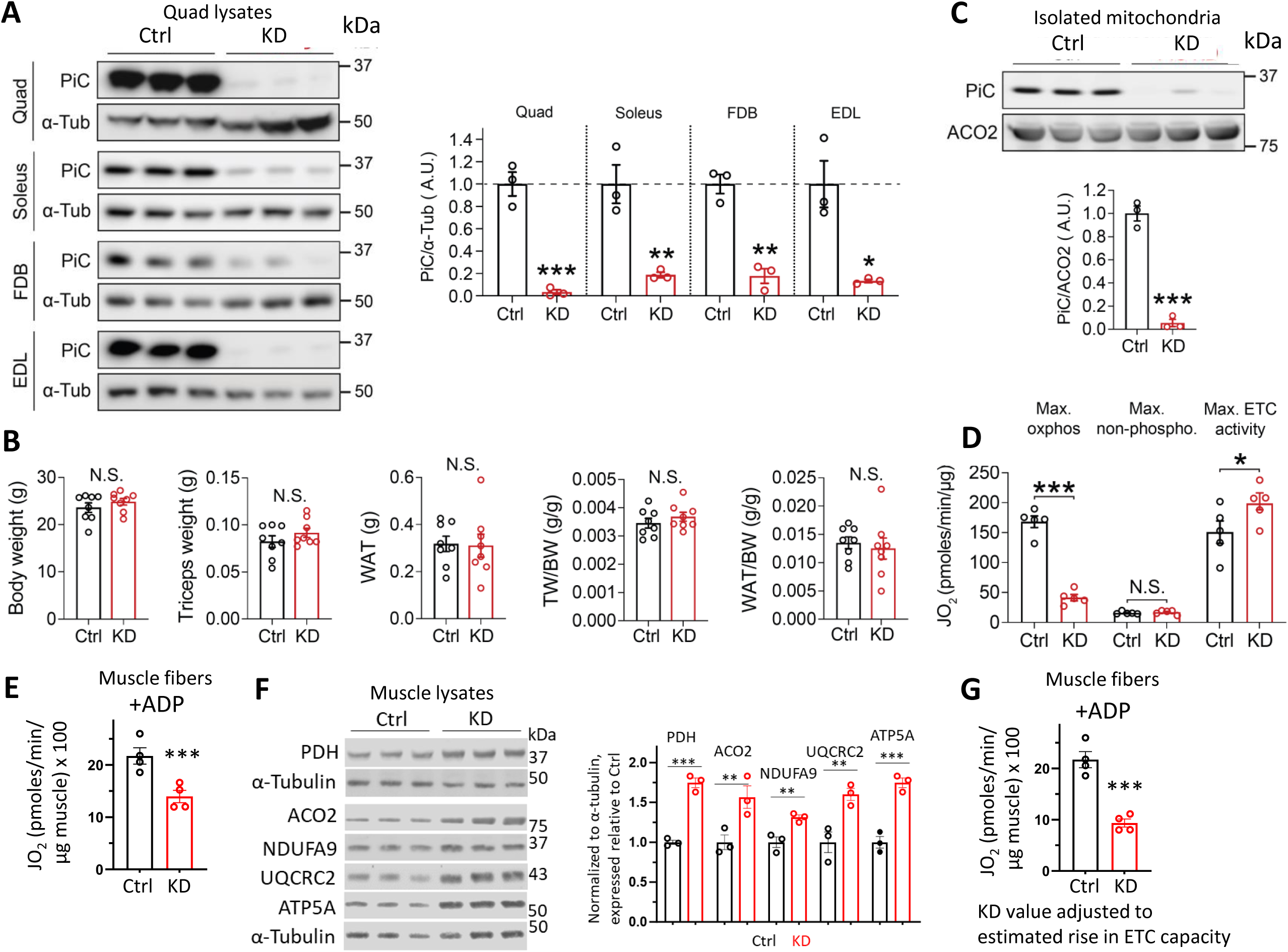
PiC depletion in SM impairs oxidative phosphorylation via Pi limitation, and is without effect on body or muscle mass in young adult mice (A) Western blots of PiC and α-Tubulin (loading control) of lysates from *Quadriceps* (Quad), *Soleus*, *Flexus digitorum longus* (FDB) and *Extensor digitorum longus* (EDL), from Ctrl and PiC KD mice. *Left*: representative Western blots, with each lane showing PiC expression in Quad from a different mouse. *Right*: Averaged values of PiC relative to α-Tubulin, then expressed as a fraction of the mean Ctrl value; n = 3/genotype. (B) Body weight (BW), triceps weight and white adipose tissue (WAT) weight in Ctrl and PiC KD mice; n = 7/genotype.(C) Representative Western blot and quantification of PiC in mitochondria isolated from SM dissected from all limbs. Each lane shows a sample from a different mouse. Aconitase 2 (ACO2) was used as the loading control; n = 3/genotype. (D) Oxygen consumption rate (JO_2_) measured in isolated SM mitochondria supplied with pyruvate/malate (P/M; 10 mM/5 mM). Max. oxphos: JO_2_ reflects maximal oxidative phosphorylation with saturating [ADP] and [substrate]. Max non-phospho: JO_2_ reflects maximal non-phosphorylating oxidation when oligomycin was used to inhibit the ATP synthase. Max ETC activity: JO_2_ reflects the maximal electron transport chain (ETC) activity when the chemical uncoupler, FCCP (1 µM), was used; n=5/genotype. (E) JO_2_ measured in permeabilized fibers from *Extensor digitorum brevis* muscle supplied with saturating P/M (10 mM/5 mM) and ADP (3 µM). Values were normalized to the wet weight of the muscle, measured after each series of JO_2_ experiments; n=5/genotype. (F) Western blot showing protein expression of major substrate oxidation and ETC proteins in Quad lysate; each lane shows data from a different mouse. Right: Quantification, n=3/genotype. PDH: pyruvate dehydrogenase (E2 subunit). NDUFA9: Complex I subunit. UQCRC2: Complex III subunit. ATP5A: ATP synthase subunit. (G) JO_2_ values from panel E, with values from KD fibers adjusted for an assumed 30% increase in substrate oxidation and ETC protein abundance, based on data from panel F. All bar charts: individual points represent values from separate mice, and bars represent mean ± s.e.m. Statistical comparison: unpaired t-test, \**p*<0.05; **, p<0.01; \*\*\**p*<0.001; N.S.: not significant.

PiC KD mice were born at a slightly higher than expected Mendelian ratio (41% Cre-, 59% Cre+; 11 litters, 93 mice), remained healthy well into adulthood, and mortality was not observed in mice as old as 1 year of age. Several models of mitochondrial dysfunction in SM resulted in myopathy characterized by muscle atrophy and loss of body weight, and early mortality (47–50). Because we aimed to evaluate mitochondrial Ca^2+^ handling and cytosolic Ca^2+^ dynamics without concurrent pathology, we evaluated KD mice for body and muscle mass, and identified 13 wks of age as a time point when these measurements, and also white adipose tissue mass, were similar in KD and Ctrl mice (Fig 1B). Unless otherwise indicated, experiments were conducted using mice at this time point.

PiC is expected to be the predominant source of Pi for ATP synthesis by oxidative phosphorylation (oxphos); thus we expected that PiC KD in SM would limit oxphos. To test this, bioenergetics analyses were performed using mitochondria isolated from all limb muscles; in KD mice, this preparation showed a ∼95% depletion of PiC (**Fig 1C**). Substrate oxidation was evaluated as O_2_ consumption rate (JO_2_) in mitochondria energized with pyruvate and malate (P/M), in the presence of 1 mM Pi. Maximal oxphos (saturating P/M and ADP) was ∼70% lower in KD mitochondria, whereas JO_2_ in the presence of oligomycin (ATP synthase inhibition; to measure maximal non-phosphorylating (leak) JO_2_), was similar between KD and Ctrl (Fig 1D). These observations are consistent with a Pi limitation on oxphos in KD mitochondria. We noted that maximal uncoupled (i.e., +FCCP) JO_2_ was significantly higher in the KD (Fig 1D), indicating a greater electron transport chain (ETC) capacity as documented in other mouse models and in cells from humans with deficient oxphos (51–55), and thus not a primary effect of PiC KD.

To further explore oxphos capacity in KD SM mitochondria, bioenergetics analyses were performed using permeabilized fibers from *Flexor digitorum longus* (FDB) muscle and saturating concentrations of pyruvate and malate, and ADP; JO_2_ in response to maximal [ADP] was 35% lower in KD fibers (Fig 1E). Evaluating the expression of major substrate and ETC proteins in Quad lysates revealed a ∼30% to 70% rise over Ctrl (Fig 1F). Assuming a conservative 30% increase in substrate plus ETC protein capacity in KD muscle, maximal oxphos in SM fibers, considered relative to substrate plus ETC protein capacity, would be ∼60% lower in KD fibers (Fig 1G), approaching maximal oxphos in KD mitochondria. Altogether these bioenergetics analyses show that KD mitochondria exhibit the expected Pi limitation of oxphos, leading to a substantial (70%) decrease in oxphos capacity, though compensation in ETC protein capacity at the whole-muscle level lessens the oxphos defect.

Other transporters in the IMM can transport Pi: the dicarboxylate carrier (DiC, encoded by SLC25A10) and three ATP-Mg^2+^-Pi transporters (SCaMC-3/-1/-2; encoded by SLC25A23/24/25 respectively) (56). To check if any of these could compensate for PiC loss, we report the LFQ values from a proteomics dataset that we obtained from isolated SM mitochondria. From this analysis we calculated a 97% decrease in PiC in KD mitochondria (**SFig 1B**), comparable to the ∼95% decrease obtained by western blot of isolated SM mitochondria (Fig 1C). All other potential Pi transpoters, except SCaMC-3, were detected and had similar expression in KD and Ctrl mitochondria. Yet, the abundance of each was two orders of magnitude lower than for PiC in Ctrl mitochondria, and, altogether, accounted for only 4% of the total theoretical Pi transport protein abundance (**SFig 1B**). Furthermore, the dicarboxylate transporter (DiC), the only other possible mitochondria Pi transporter with a transport rate approaching that of the PiC (see Discussion), was only 0.5% of the theoretical Pi transporter protein abundance in SM mitochondria from Ctrl mice. Thus the PiC is likely the only relevant Pi transporter in Ctrl and KD mitochondria in mouse SM.

### Increased mitochondrial Ca^2+^ uptake and [Ca^2+^]m in PiC-depleted mitochondria

Pi uptake provides a source of weak acid that would enable ΔΨm-driven Ca^2+^ uptake into mitochondria. In the matrix, Pi would bind Ca^2+^, thereby decreasing the concentration of free Ca^2+^ in the matrix . We tested how PiC depletion would impact these processed by evaluating Ca^2+^ uptake and matrix free [Ca^2+^] ([Ca^2+^]m) in isolated KD and Ctrl SM mitochondria. To directly compare the amount of Ca^2+^ uptake by mitochondria and the corresponding free Ca^2+^ change in the matrix, we set up simultaneous measurements of the Ca^2+^ concentration in the suspension buffer ([Ca^2+^]c), using rhodFF, and [Ca^2+^]m, using FuraFF compartmentalized to the mitochondrial matrix (see Methods). CGP37157 was included, to inhibit Ca^2+^ efflux, and thapsigargin, to inhibit Ca^2+^ uptake by any sarcoplasmic reticulum stores copurified with mitochondria. Additionally, mitochondria were first depleted of endogenous Pi by incubation with glucose, hexokinase and ADP, pelleting, then resuspending in incubation medium devoid of these components (32). Unless otherwise stated, 1 mM Pi was included in the incubation medium.

Using mitochondria energized with pyruvate and malate, [Ca^2+^]c was monitored after a bolus addition of 10 µM CaCl_2_ that increased [Ca^2+^]c to ∼4 μM (Fig 2AE). CaCl_2_ clearance was Ruthenium Red (RuR)-sensitive in both genotypes, confirming that it was mtCU-mediated (Fig 2A middle). PiC KD mitochondria not only could take up Ca^2+^, the amount of Ca^2+^ cleared after 30 seconds was ∼40% greater in KD vs. Ctrl (Fig 2A). To test if PiC deletion affects Ca^2+^ uptake by suppressing Pi uptake, we measured Ca^2+^ uptake in zero Pi which clearly decreased Ca^2+^ uptake (by ∼30%) in the Ctrl but had less of a suppressing effect (∼16%) in the PiC KD (Fig 2B). Thus, SM mitochondria displayed some Pi-dependence of Ca^2+^ uptake, as documented in mitochondria from rat brain (32) and guinea pig heart (33).

**Figure 2:**
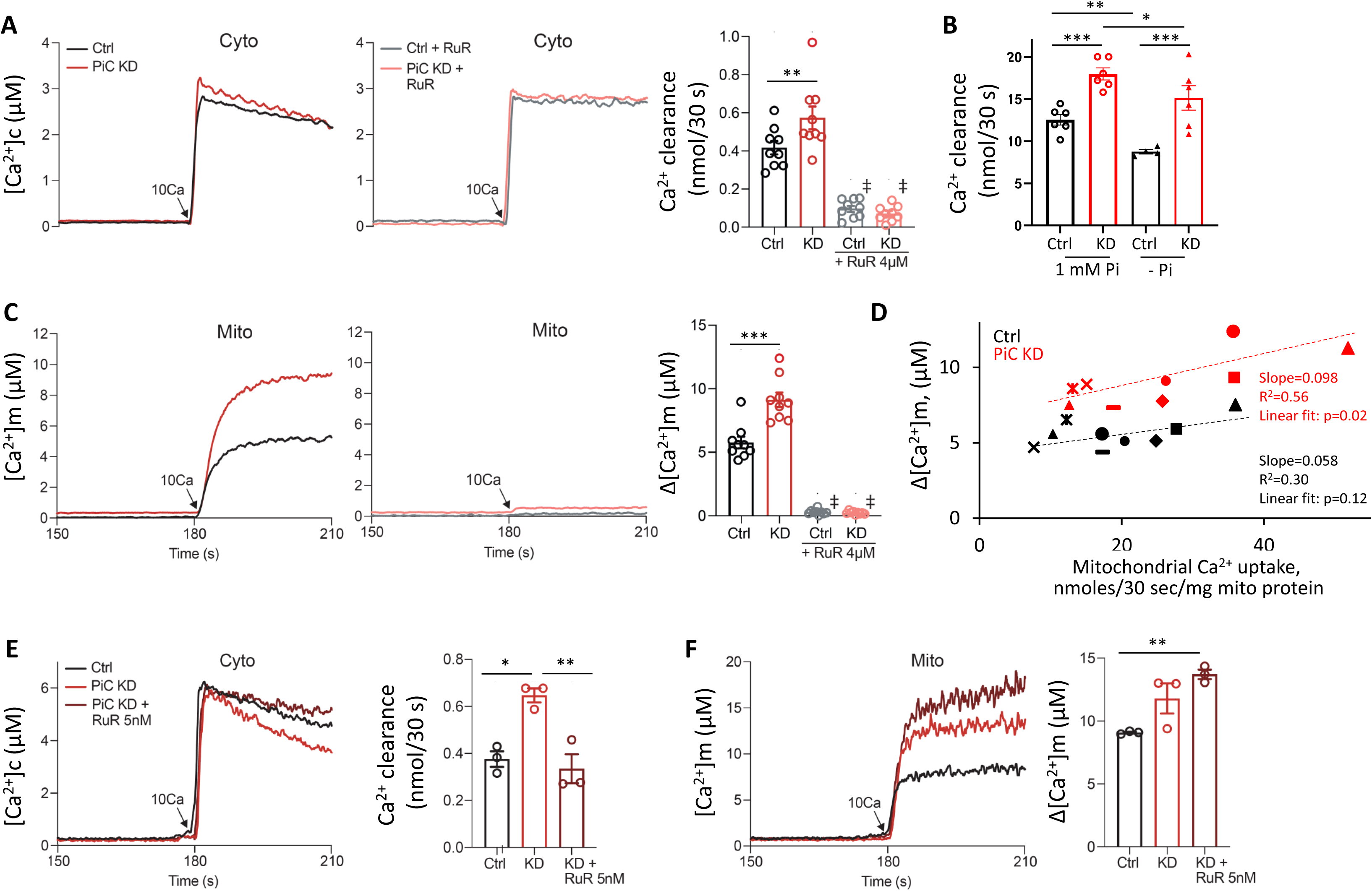
Ca2+ uptake and [Ca2+]m are elevated in PiC-depleted SM mitochondria. (A) *Left:* Representative time course of the mitochondrial clearance from the buffer (“Cyto”) of a 10 µM CaCl_2_ bolus (10Ca) by suspensions of Ctrl (black) and PiC KD (red) mitochondria. *Middle*: The same experiment was done in the presence of Ruthenium Red (RuR; 4 µM) to evaluate specificity of Ca^2+^ clearance by the mtCU (Ctrl: gray; PiC KD: light red). *Right*: Quantification of mitochondrial Ca^2+^ clearance, measured as the change in concentration of Ca^2+^ in the buffer, [Ca^2+^]c, over 30 seconds; n = 9/experimental condition/genotype. (B) Ca^2+^ clearance measured in isolated mitochondria after a 10 µM CaCl_2_ bolus addition, in the presence and absence of 1 mM Pi; n = 4-6/experimental condition. Clearance rate was calculated as the first derivative of an exponential function that was fit to the data. (C) Representative time course of the concentration of free Ca^2+^ in the mitochondrial matrix ([Ca^2+^]_m_) upon addition of a 10 µM CaCl_2_ bolus (10Ca) in suspensions of Ctrl (black) and PiC KD (red) mitochondria. *Middle*: The experiment was repeated in the presence of RuR (4 µM; middle panel), in Ctrl (gray) and PiC KD (light red). *Right*: Quantification of the [Ca^2+^]_m_ 30 s after the CaCl_2_ bolus; n = 9/experimental condition. (D) Average values of [Ca^2+^]m and Ca^2+^ uptake, from each mitochondrial isolation (i.e., from each mouse), in response to a 10 µM CaCl_2_ bolus were plotted. Different symbols were used for different experimental days. Black symbols: Ctrl mitochondria; red symbols: PiC-depleted mitochondria. Lines: data within a genotype were fitted by a linear regression model; R^2^: goodness of fit of the model. Data are from experiments shown in panels A and C. (E) Representative time course of the mitochondrial clearance of a 10 µM CaCl_2_ bolus (10Ca) in suspensions of Ctrl (black), PiC KD (red) mitochondria, and in PiC KD mitochondria in the presence of 5 nM RuR to partially inhibit mtCU (dark red). *Right*: Quantification of the mitochondrial Ca^2+^ uptake, measured as in panel A; n = 3/ experimental condition. (F) Representative time course of the mitochondrial matrix Ca^2+^ ([Ca^2+^]_m_) upon addition of a 10 µM CaCl_2_ bolus (10Ca) in suspensions of Ctrl (black), PiC KD (red) mitochondria and in PiC KD mitochondria in presence of 5 nM RuR (dark red). *Right*: Quantification of the [Ca^2+^]_m_, 30 s after the CaCl_2_ bolus (n = 3/experimental condition). All bar charts: individual data are values from separate mice, and bars represent mean ± s.e.m. Statistical comparison: two-way ANOVA (Panel A-C) or one-way ANOVA (Panel D, E); p values are from post hoc tests (Bonferroni corrections for multiple comparisons) \**p*<0.05; \*\**p*<0.01; \*\*\**p*<0.001; and # *p*<0.001 is vs. no RuR (same genotype).

Measurement of [Ca^2+^]m, simultaneously with [Ca^2+^]c, allowed direct comparison of both parameters in each preparation. In response to 10 µM CaCl_2_ addition, [Ca^2+^]m rose; most of the increase occurred within 10 seconds of CaCl_2_ addition, and was followed by a phase of slower uptake (Fig 2C left). The rise in [Ca^2+^]m was greater in KD mitochondria; after 30 seconds, it was ∼60% higher than in PiC-replete mitochondria (Fig 2C right). The rise in [Ca^2+^]m in both genotypes was RuR-sensitive and thus mtCU-mediated (Fig 2C middle, right). To test if the greater rise in [Ca^2+^]m in KD mitochondria might simply be due to the greater uptake of Ca^2+^, we minimized the difference in Ca^2+^ clearance between KD and Ctrl by adding a low concentration of RuR (5 nM) to KD mitochondria (Fig 2E); [Ca^2+^]m remained substantially higher in KD vs. Ctrl, even when uptake was similar in the KD and Ctrl (Fig 2F).

To further analyze the impact of PiC depletion on [Ca^2+^]m as a function of Ca^2+^ uptake, we took advantage of some day-to-day variability. For each isolated mitochondria preparation, the mean [Ca^2+^]m value was plotted against the corresponding mean Ca^2+^ uptake value (Fig 2D: each symbol represents a different experimental day, and black symbols are from the Ctrl and red from the KD preparation). This analysis showed that, in Ctrl mitochondria, there was little relationship between [Ca^2+^]m and Ca^2+^ uptake (p=0.12 for the linear regression model), while, in PiC KD mitochondria, all [Ca^2+^]m values are shifted above the Ctrl [Ca^2+^]m values and their dependence on Ca^2+^ uptake was significant (p=0.02; Fig 2D).

### PiC KD mitochondria depolarize more during Ca^2+^ uptake

Pi uptake by the PiC is hypothesized to mitigate ΔΨm loss during mitochondrial Ca^2+^ uptake. To this end, ΔΨ_m_ was evaluated using TMRM. Energizing mitochondria with pyruvate and malate prior to CaCl_2_ addition polarized ΔΨ_m_ to the same extent in Ctrl and KD (Fig 3A)_._ In response to a 10 µM CaCl_2_ bolus, some depolarization was noted in both genotypes, but the magnitude was consistently greater in KD mitochondria (**Fig3A**). Though this greater ΔΨ_m_ response in PiC-depleted mitochondria might reflect matrix alkalinization resulting from Pi lack, other mechanisms were possible. In particular, we asked whether any residual ADP in the matrix combined with the greater matrix free Ca^2+^ in KD mitochondria and oxphos capacity could lead to a greater stimulation of oxphos in KD than in Ctrl mitochondria and consequently a greater depolarization in KD mitochondria. This was tested by measuring ΔΨ_m_ during the CaCl_2_ bolus in the presence of oligomycin to inhibit the ATP synthase. Oligomycin addition caused both KD and Ctrl mitochondria to hyperpolarize (**Fig3B**), indicating that some oxphos occurred in the absence of oligomycin in both the Ctrl and KD. Addition of 10 µM CaCl_2_ caused both KD and Ctrl mitochondria to depolarize, again to a greater extent in KD mitochondria (**Fig3B**); thus the larger depolarization in the KD cannot be explained by a greater Ca^2+^-dependent stimulation of oxphos.

**Figure 3:**
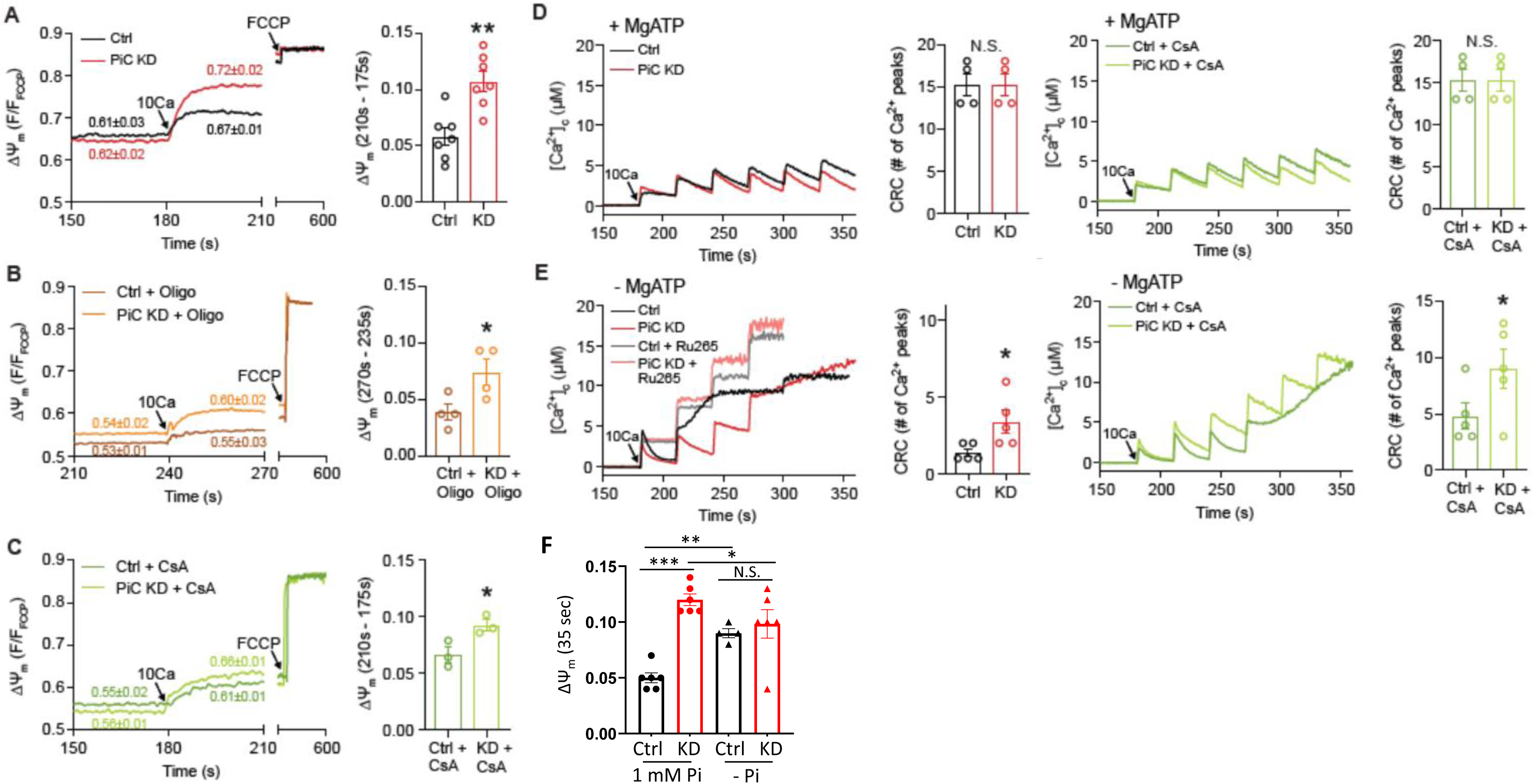
PiC-depleted SM mitochondria depolarize more in response to a Ca2+ pulse. (A) Mitochondrial membrane potential (ΔΨ_m_) was measured using TMRM in suspensions of Ctrl (black) and PiC KD (red) mitochondria. TMRM was used in de-quench mode, therefore the direction of membrane polarization is downward. FCCP was used to completely dissipate the ΔΨ_m_. *Right*: Quantification of ΔΨ_m_ 30 s after a 10 µM CaCl_2_ bolus; n = 7/genotype. (B) ÄØ_m_ measurements in suspensions of Ctrl (brown) and PiC KD (orange) mitochondria, in the presence of oligomycin (Oligo; 2.5 µg/µl) to inhibit the ATP synthase. *Right*: Quantification of ΔΨ_m_ 30 s after a 10 µM CaCl_2_ bolus; n = 4/genotype. (C) ÄØ_m_ measurements in suspensions of Ctrl (green) and PiC KD (light green) mitochondria, in the presence of cyclosporin A (CsA; 5 µM) to decrease permeability transition pore (PTP) open probability. *Right*: Quantification of ΔΨ_m_ 30 s after a 10 µM CaCl_2_ bolus; n = 3/genotype. (D) Evaluation of Ca^2+^ overload-induced mitochondrial PTP opening in Ctrl (black) and PiC KD (red) mitochondria suspended in medium containing Mg^2+^-ATP, without CsA (*left*) and with CsA (*right*) (with CsA: Ctrl in dark green, PiC KD in light green). [Ca^2+^]_c_ was recorded during repeated addition (every 30 s) of 10 µM CaCl_2_ boluses. *Right*: Quantification of the Ca^2+^ retention capacity (CRC); n = 4/genotype. (E) Evaluation of Ca^2+^ overload-induced mitochondrial PTP opening in Ctrl (black) and PiC KD (red) mitochondria suspended in medium devoid of Mg^2+^-ATP, without CsA (*left*) and with CsA (*right*) (with CsA: Ctrl in dark green, PiC KD in light green. [Ca^2+^]_c_ was recorded during repeated addition (every 30 s) of 10 µM CaCl_2_ boluses. Some experiments were performed in the presence of Ru265 to inhibit the mtCU (Ctrl, light gray and PiC, light red). *Right*: Quantification of the Ca^2+^ retention capacity (CRC); n = 5/condition/genotype. (F) Measurements of ΔΨ_m_ in isolated mitochondria (as in Panel A) after a 10 µM CaCl_2_ bolus addition, in the presence and absence of 1 mM Pi; n = 4-6/condition/genotype. All bar charts: individual data points are from separate mice, and bars represent mean ± s.e.m. Statistical comparison: Panels A-E, unpaired t-test, \**p*<0.05; \*\**p*<0.01; N.S.: not significant. Panel F, two-way ANOVA, p values are from post hoc tests (Bonferroni corrections for multiple comparisons) \**p*<0.05; \*\**p*<0.01; \*\*\**p*<0.005; N.S.: not significant.

Considering that PiC and/or P_i_ may regulate the permeability transition pore (PTP) (57–60), we also tested if the greater Ca^2+^-dependent depolarization in KD mitochondria was related to PTP opening. Measurements of ΔΨ_m_ in response to 10 µM CaCl_2_ were repeated in the presence of the PTP inhibitor cyclosporin A (CsA); the KD still exhibited a greater depolarization in the presence of CsA (**Fig3C**). We further compared the ability for the PTP to open in KD *vs*. Ctrl preparations by evaluating the Ca^2+^ retention capacity in response to a series of 10 µM CaCl_2_ pulses. Ca^2+^ clearance was still evident after ∼ 15 pulses in both Ctrl and KD mitochondria before FCCP was added to fully depolarize mitochondria (**Fig3D**). Thus, PTP open probability was low under the conditions used for Ca^2+^ and ΔΨ_m_ measurements, likely because of the inclusion of adenine nucleotides (ATP-Mg^2+^) in the incubation medium (for review: (61)). On the other hand, mitochondria incubated in medium devoid of ATP-Mg^2+^ and exposed to 10 µM CaCl_2_ pulses robustly underwent PTP opening, evidenced by an abrupt and substantial increase in [Ca^2+^]c and loss of ΔΨ_m_ (**Fig3E**). Fewer pulses were needed for PTP opening in Ctrl compared to PiC-depleted mitochondria (**Fig3E**), as shown previously in heart mitochondria depleted of PiC by >90% (41). PTP opening could be inhibited by CsA, and CsA sensitivity appeared similar in the Ctrl and KD (**Fig3E**), even though protein expression of cyclophilin D (CypD), which increases PTP open probability (61), was higher in KD mitochondria by ∼35% (**SFig2**). Altogether, this analysis of PTP opening suggests that PiC KD mitochondria *in situ* would be more resistant to PTP activation, despite higher CypD expression. However PTP opening was minimal in the conditions used to measure Ca^2+^ and ΔΨ_m_ and thus would not explain the greater depolarization in KD mitochondria exposed to CaCl_2_.

Finally, we checked the Pi-dependence of the CaCl_2_-induced depolarization. In the absence of Pi in the incubation medium, the depolarization elicited by 10 µM CaCl_2_ was similar in Ctrl and KD mitochondria, and the extent of the depolarization was similar to that elicited by CaCl_2_ in KD mitochondria incubated with 1 mM Pi (**Fig3F**).

### Higher protein expression of mtCU components in PiC-depleted mitochondria

Since PiC KD mitochondria exhibited greater RuR-inhibitable Ca^2+^ uptake, and this cannot be attributed to an increased driving force for Ca^2+^ uptake, we questioned whether PiC-depleted mitochondria had a greater mtCU protein abundance. Western blot analysis to evaluate the expression of the individual mtCU components in isolated SM mitochondria revealed that protein levels of MCU, EMRE and MICU1 were elevated by ∼50% in KD mitochondria when compared to 2 other mitochondria proteins, aconitase (ACO2) and TOMM20 (Fig 4A).

**Figure 4:**
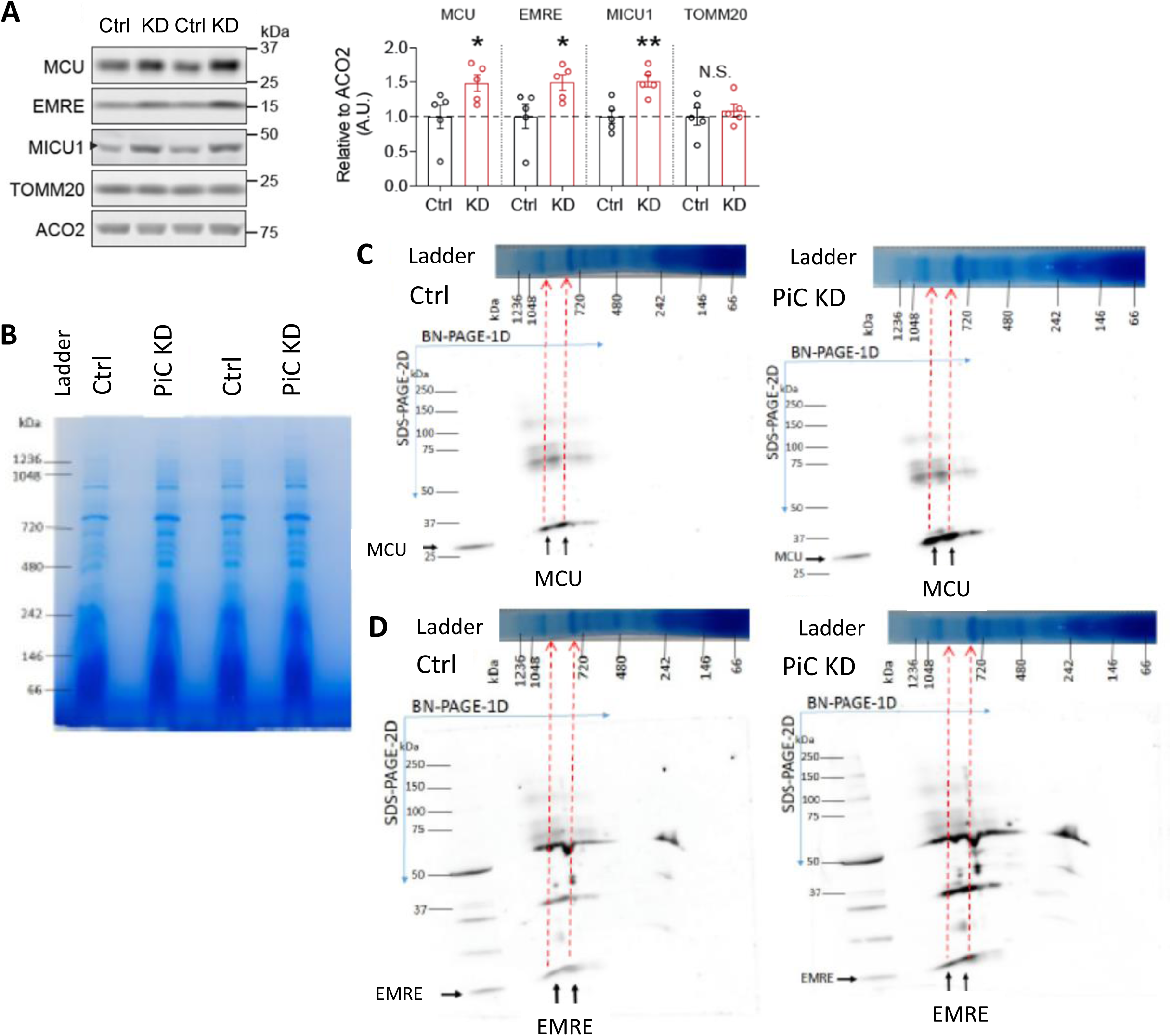
Elevated protein expression of mtCU components and assempled mtCU in PiC KD mitochondria. (A) Representative Western blots of MCU, EMRE, MICU1, TOMM20, and ACO2 in mitochondria isolated from SM from Ctrl and PiC KD mice. *Right*: Quantification relative to ACO2 (as a non-mtCU protein); n = 5/genotype. Bar charts: points are from different mice, and bars represent mean ± s.e.m. Statistics: unpaired t-test, \**p*<0.05; \*\**p*<0.01; N.S.: not significant. (B) Skeletal muscle mitochondria isolated from 2 Ctrl and 2 PiC KD mice were electrophoresed under native conditions (BN-PAGE: blue native PAGE). Shown is the Coomassie blue-stained gel. These samples were used for 2^nd^ dimension electrophoresis under denaturing conditions (panels C, D). (C) , **(D)** Lanes were cut from the native gel (shown intact in panel B, and cut and rotated 90° in panels C and D; BN-PAGE, 1^st^ dimension (1D), molecular weights of the native proteins are shown. Samples were electrophoresed under denaturing conditions (SDS-PAGE 2^nd^ dimension, 2D), transferred onto nitrocellulose membrane, then probed with antibodies against MCU **(C)** and EMRE **(D)**. Dashed lines with arrows point to the approximate molecular weight on the native gel from which the immunoreactive bands originated. As a positive control for the antibodies, skeletal muscle mitochondria prepared in loading buffer for Western blotting were run in the leftmost lane; arrows show the immunoreactive bands for this sample.

To evaluate if abundance of the mtCU complex was also increased, isolated mitochondria were electrophoresed under non-denaturing conditions, followed by a 2^nd^ dimension (2D) gel run under denaturing conditions and probed with antibodies against MCU and EMRE (**Fig4B-D**). In Ctrl mitochondria, MCU and EMRE immunoreactive bands were detected, and corresponded to complexes of ∼750 kDa and ∼1000 kDa molecular weight (MW) in the native gel (**Fig4CD**), similar to what was observed for MCU expression in cortical neurons (62) and heart mitochondria (63). The KD samples showed the same pattern as the Ctrl, but the abundance of the MCU- and EMRE-immunoreative bands was higher (**Fig4CD**). The Coomassie-stained gel (Fig 4B), and Complex II subunit SDHA (**SFig3A**), served as loading controls.

To estimate mtCU abundance at the whole muscle level, we evaluated MCU expression by Western blotting in Quad lysates. Relative to α-tubulin, MCU protein was ∼2.2x higher in KD *vs*. Ctrl (**SFig3B**). For comparison, mitochondrial proteins from the outer mitochondrial membrane (TOMM20), and matrix (ACO2, TFAM, CypD) were also detected; each was increased by ∼1.5-2x in KD SM, supporting a generalized increase in the abundance of mitochondria in KD SM, but disproportionate increase in MCU (**SFig3B; see also** Fig 5I).

**Figure 5:**
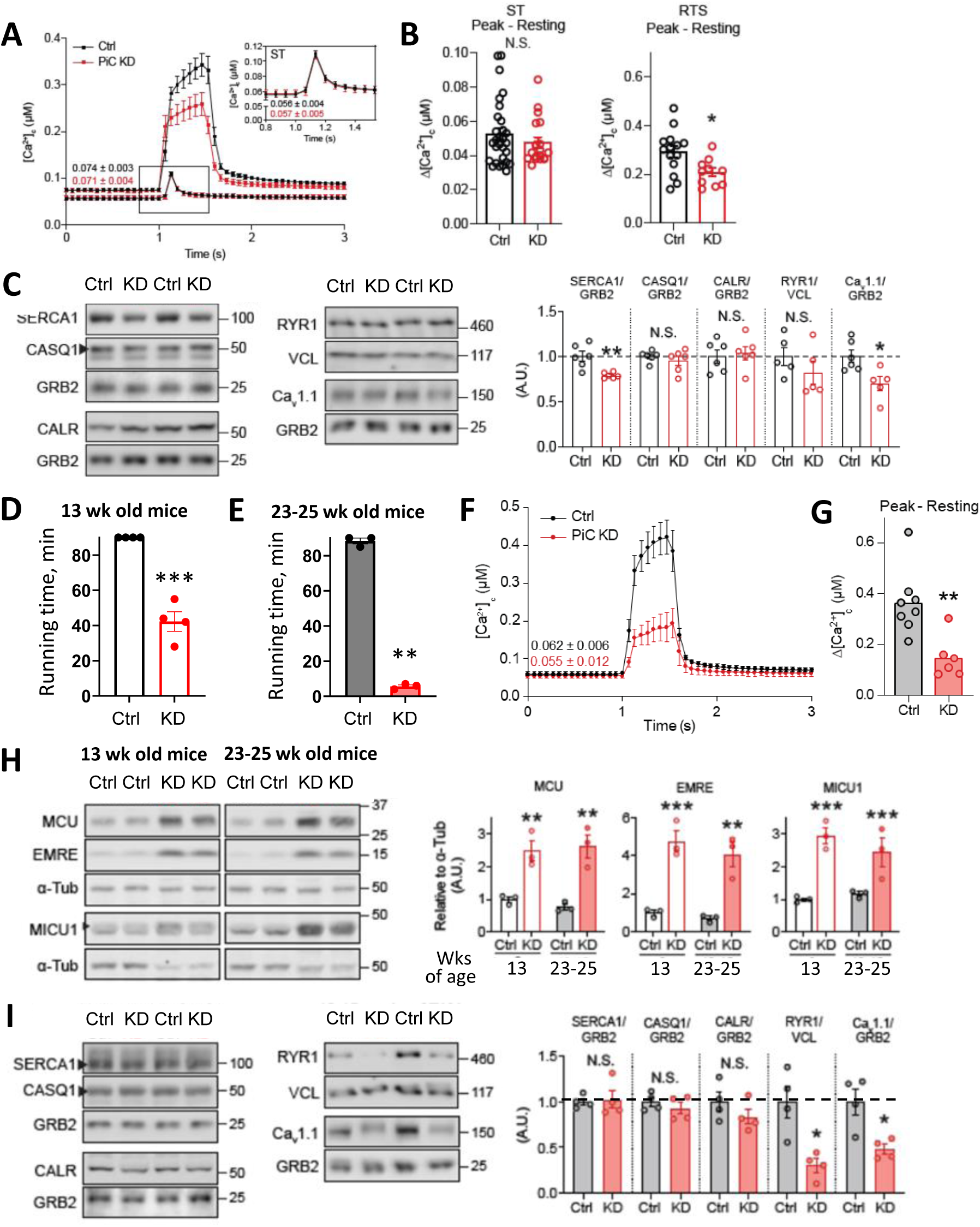
Greater exercise intolerance and suppression of the cytoplasmic Ca2+ response to tetanic muscle stimulation in PiC KD mice. (A) Traces of average (± s.e.m.) cytoplasmic Ca^2+^ concentration ([Ca^2+^]_c_) levels during single twitch (ST) and repetitive tetanic stimulation (RTS) in Ctrl (black) and PiC KD (red) *Flexor digitorum brevis* (FDB) muscle from 13-wk-old mice. Values of resting [Ca^2+^]_c_ prior to RTS are shown above the traces. *Inset*: Amplified view of averaged traces of [Ca^2+^]_c_ levels during ST, with values of resting [Ca^2+^]_c_ shown below the traces. (B) Quantification of data from (B): peak [Ca^2+^]_c_ during ST (n = 29 Ctrl and 19 PiC KD FDB), and during RTS (n = 14 Ctrl and 10 PiC KD FDB). (C) Representative Western blots from *Quadriceps* lysates, from 13-wk-old mice, of SERCA1, calsequestrin (CASQ1), calreticulin (CALR), Type I ryanodine receptor (RYR1) and Cav1.1 (L-type Ca^2+^ channel, 1S subunit). GRB2 and vinculin (VCL): loading controls. *Right*: Mean values relative to GRB2 (n = 6/genotype) or VCL, n = 5/genotype. (D) Treadmill running time to exhaustion by 13-wk-old mice, n=4/genotype. (E) Treadmill running time to exhaustion by 23-25-wk-old mice, n=4/genotype. (F) Traces showing the average (± s.e.m.) cytoplasmic Ca^2+^ concentration ([Ca^2+^]_c_) during repetitive tetanic stimulation (RTS) in *Flexor digitorum brevis* (FDB) muscle from 23-25-wk-old Ctrl (black) and PiC KD (red) mice. Values of resting [Ca^2+^]_c_ are shown above the traces. (G) Quantification of data from (F): peak [Ca^2+^]_c_ during RTS in FDB, n=8 FDB fibers/genotype. (H) Representative Western blots of mitochondrial Ca^2+^ uniporter components in *Quadriceps* lysates from 13-wk-old and 23-25-wk-old mice. *Right*: Averaged values relative to α-Tubulin; n = 3/genotype. (I) Representative Western blots from *Quadriceps* lysates, from 23-25-wk-old mice, of SERCA1, calsequestrin (CASQ1), calreticulin (CALR), Type I ryanodine receptor (RYR1) and Cav1.1 (L-type Ca^2+^ channel, 1S subunit). GRB2 and vinculin (VCL) were used as a loading controls. *Right*: Averaged values relative to GRB2 (n = 4/genotype) or VCL; n = 4/genotype. Bar charts: individual data points are from separate mice or fibers, and bars represent mean ± s.e.m. Statistical comparison: unpaired t-test, \**p*<0.05; \*\**p*<0.01, *** *p*<0.001; N.S.: not significant.

### Deteriorating cytoplasmic Ca^2+^ response in KD SM parallels progressive exercise intolerance

Considering the cross-talk between Ca^2+^ uptake by SM mitochondria and Ca^2+^ release from the sarcoplasmic reticulum (SR) (64–66), we evaluated the impact of PiC KD on [Ca^2+^]c transients in SM fibers. [Ca^2+^]c responses were evaluated in isolated *Flexus digitorum brevis* (FDB) fibers during single twitch (ST) and repeated tetanic stimulation (RTS), done sequentially in each FDB preparation. A similar pre-stimulus baseline [Ca^2+^]_c_ of ∼50-60 nM was observed in KD and Ctrl fibers (Fig 5A). In both genotypes, ST stimulation evoked a ∼50 nM cytoplasmic Ca^2+^ transient (**Fig5AB**) that decayed with a similar kinetic (**SFig4**), then returned to a similar baseline level (**Fig5A**). Ctrl fibers challenged with RTS displayed a rapid-onset, higher amplitude rise in [Ca^2+^]c, initially of ∼300 nM, and that continued to increase with a slower kinetic throughout the stimulation period (**Fig5AB**). Notably, the local [Ca^2+^]c rise sensed by mitochondria close to SR Ca^2+^ release exceeded the global [Ca^2+^]c rise we measured, and likely reached several micromolar (66). KD fibers challenged with RTS responded with the same pattern of cytoplasmic Ca^2+^ elevation as Ctrl fibers, but with two differences. First, the initial rapid phase tended to be faster in KD fibers (**Fig5A**). Second, the amplitude of the [Ca^2+^]c rise was less throughout the stimulation period, by ∼30% (**Fig5AB**). These differences in KD fibers were unrelated to major changes in the abundance of major extra- mitochondrial Ca^2+^ handling proteins, including SR/ER Ca^2+^ release or uptake channels (type 1 ryanodine receptor (RyR1), SERCA1), Ca^2+^ binding proteins (calsequestrin (CASQ1), calreticulin (CALR)), or to the major SM subunit of the L-type Ca^2+^ channel (Cav1.1) (**Fig5C**).

As integrative functional readouts of Ca^2+^- and also ATP-dependent processes, we tested exercise capacity by treadmill running, and SM function by ex vivo force measurements in stimulated *Extensor digitorum longus* (EDL). PiC-depleted mice were exercise intolerant, evidenced by a ∼25% decrease in running time on a treadmill (**Fig5D**). Isolated KD EDL, which had similar cross-sectional area to the Ctrl (**SFig5A**), responded to a ST with only minor differences from Ctrl in peak force generation and the time to attain this force (**SFig5BC**). Challenged with RTS, KD muscle again generated a similar peak force as Ctrl (**SFig5D**), but took 2x longer to reach peak it (**SFig5E**), consistent with the lower [Ca^2+^]c response to a tetani in KD muscle (**Fig5C**). Finally, Ctrl and KD showed similar kinetics of force recovery from a fatigue protocol (**SFig5FG**).

Phenotypes related to major mitochondrial disruption often change with time (e.g. (41, 47–50, 52)). Thus we repeated treadmill running and cytoplasmic [Ca^2+^] responses in older PiC KD and Ctrl mice. Ctrl mice age 23-25 wks ran run as much as the 13 wk old ones (**Fig5E**). However, KD mice could run for only ∼5 mins (**Fig5E**), a dramatically lower exercise capacity compared to same-age Ctrl mice and to younger KD mice. Challenged with RTS, fibers from 23-25-wk-old Ctrl mice showed a similar pattern of response as younger Ctrl and KD mice (**Fig5FG**), but with a more profound suppression of the maximal response, which was 70% lower vs. same-age Ctrl (**Fig5FG**). Worsened exercise performance and maximal Ca^2+^ response to RTS in older KD mice occurred without further deterioration in bioenergetics capacity (**SFig6**), change in abundance of mtCU components, which remained elevated as in younger KD mice (**Fig5H**), or in levels of ER Ca^2+^ binding proteins, SERCA1 and Cav1.1, which remained similar in Ctrl and KD SM. Differently from younger KD SM, older KD SM showed substantially lower RyR1 and Cav1.1 protein levels, which were ∼70% and 50% less, respectively (**Fig5I**).

### Lowering the expression of MCU in PiC KD muscle worsens exercise capacity in younger KD mice

The foregoing observations suggest that the exercise intolerance of PiC KD mice may be causally related to their mitochondrial Ca^2+^ phenotype. The elevated mitochondrial Ca^2+^ uptake could impact exercise capacity by curtailing the cytoplasmic Ca^2+^ response to stimulation (via decreased positive feedback by Ca^2+^ on RYR1). Higher [Ca^2+^]m in KD mitochondria could also influence exercise capacity, e.g., by boosting Ca^2+^-stimulated matrix metabolism. To test these possibilities, we depleted MCU in the SM of PiC KD mice. Younger KD mice (13-wk-old) were targeted because they expressed sufficient exercise capacity to allow improvement or deterioration to be detected. Furthermore, younger KD showed little extra-mitochondrial Ca^2+^ phenotype. MCU was depleted in SM by crossing PiC^fl/fl^ mice expressing HSA-Cre with MCU^fl/fl^ mice (Cre+). MCU KD was tamoxifen (Tam) inducible (Fig 6A, left panel). Ctrl mice expressed floxed alleles for both PiC and MCU but did not express Cre (Cre-). Nine-week-old mice were treated with Tam (Tam+) or corn oil vehicle (Tam-) for 5 consecutive days. Measurements were made in 13-wk-old mice. Tam treatment of Cre+ mice lowered MCU in Quad lysates by ∼85% from the level in Cre+Tam-Quad (i.e., PiC KD only) (**Fig6A**), or by ∼25% relative to the level of MCU in Ctrl Quad (Fig 6A, right panel). Exercise capacity was tested as in **Fig5DE**, and blood lactate was also measured immediately before and after running. Cre+Tam-(PiC KD only) mice showed a similar exercise deficit as described in Fig5D (**Fig6B**, closed symbols), and this was accompanied by elevated blood lactate (**Fig6C**). PiC KD mice with lowered MCU (Cre+Tam+) ran for less time than PiC KD mice (**Fig6B**), and the rise in circulating lactate was greater (**Fig6C**). Thus the MCU increase served as an adaptation to the PiC KD-induced SM dysfunction.

**Figure 6:**
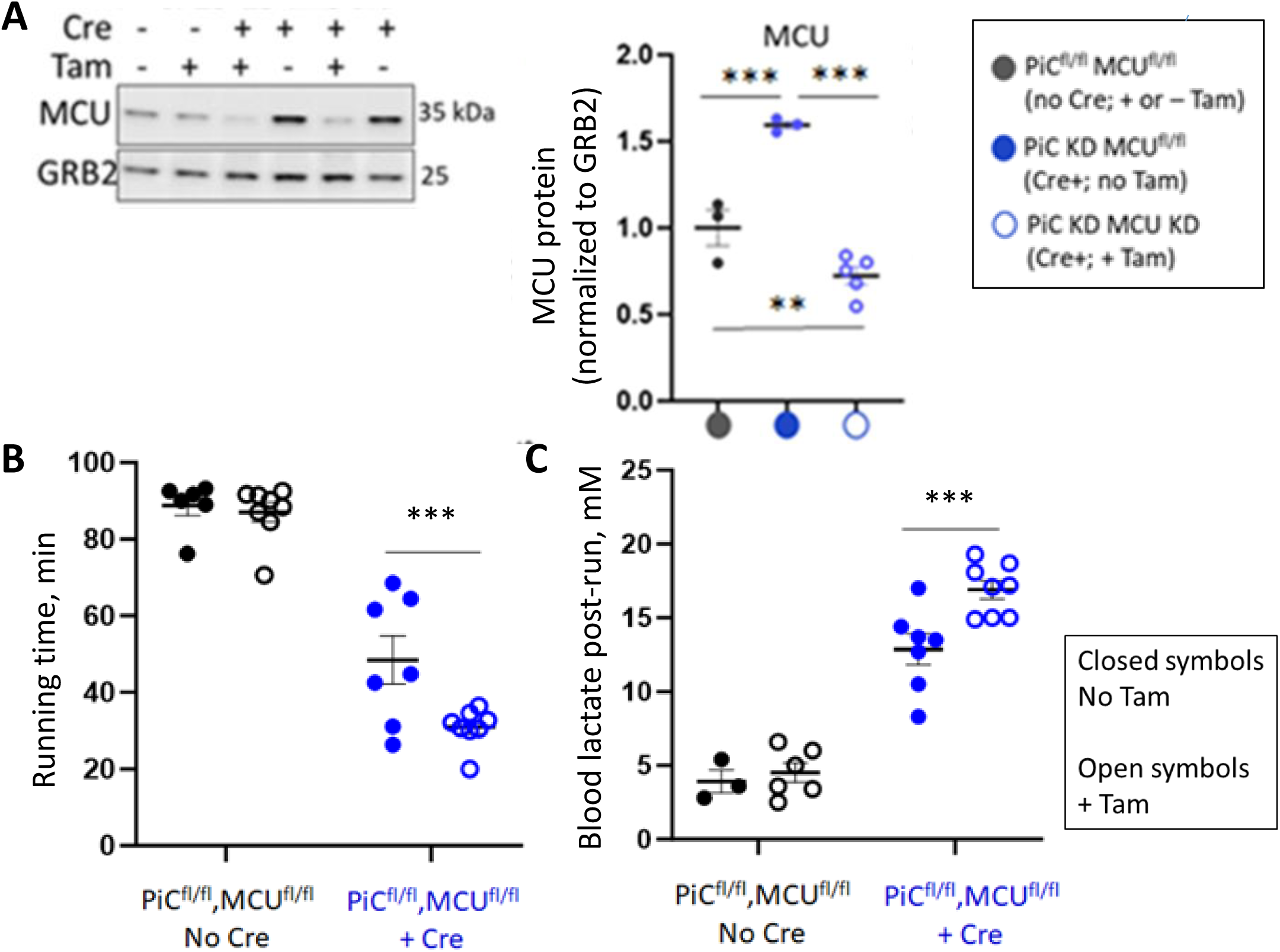
MCU decrease in PIC KD muscle worsens exercise capacity. **(A)** Representative Western blots showing Tam-dependent decrease in MCU in Quad lysates from PiC^fl/fl^ x MCU^fl/fl^ – HSA-Cre (Cre+) either treated with tamoxifen (Tam+) or not (Tam-). Ctrl mice are PiC^fl/fl^ x MCU^fl/fl^ without Cre expression (Cre-). Note that all Cre+ mice have PiC KD in SM (e.g., see Fig1). Right panel: quantification and statistical analysis: one-way ANOVA, post-hoc Tukey tests, \*\**p*<0.01, \*\*\**p*<0.01. **(B)** Treadmill running to exhaustion. Statistical analysis: two-way ANOVA, post-hoc Tukey tests, \*\*\**p*<0.01. **(C)** Blood lactate immediately before and after treadmill running. Statistical analysis: two-way ANOVA, post-hoc Tukey tests, \*\*\**p*<0.01. All panels: individual data points are from different mice, and lines and error bars represent mean ± s.e.m.

### Elevated mtCU abundance is not unique to PiC loss in SM

We noted in previous publications that Drp1 depletion in SM was accompanied by increased MCU protein and greater mitochondrial Ca^2+^ uptake (50, 67), suggesting that the increased protein abundance of mtCU components that we observed (**Fig4, 5H**) is not unique to PiC depletion. In support, using a model of whole-body Frataxin KD with a mild bioenergetics phenotype in SM (68), we found that all mtCU components were upregulated in SM mitochondria, without an increase in other mitochondrial proteins (Fig 7A) or the abundance of mitochondria determined by electron microscopy (68). Furthermore, additional analysis of a proteomics dataset from heart mitochondria (69) revealed a rise in MCU and MICU1 protein levels, and these increases were among the greatest magnitude increase in mitochondrial protein expression in each of the five models of mitochondrial dysfunction used in that study (**Fig7B**). Finally, as reported in Drp1-depleted SM (50), the higher protein levels of mtCU components did not reflect greater mRNA expression (**SFig7**). These findings suggest PiC KD shares adaptive mtCU upregulation with various impairments of mitochondrial metabolism and dynamics.

**Figure 7:**
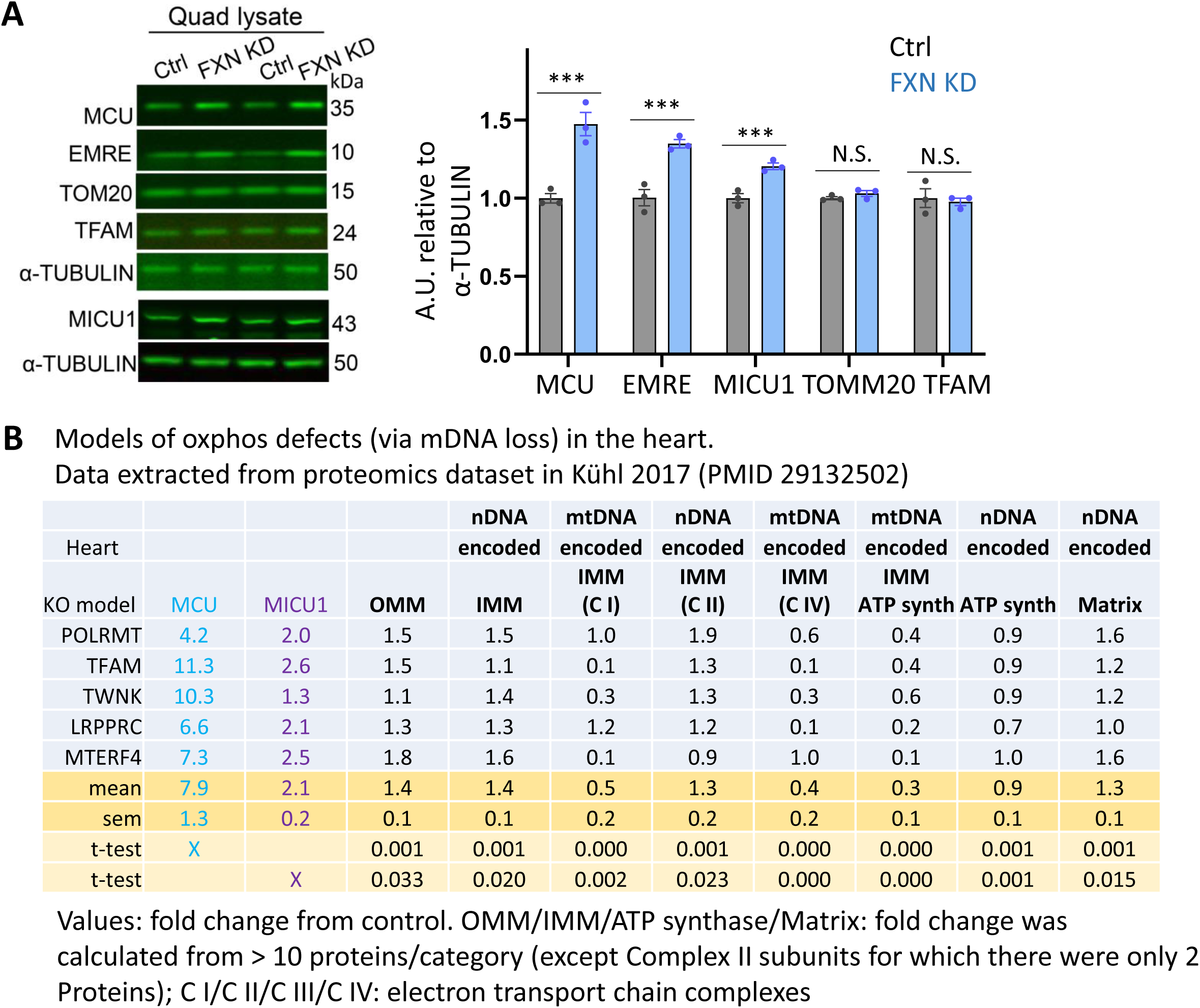
Six additional, independent models of oxphos deficiency in striated muscle showing an increased abundance of mtCU proteins. **(A)** Western blot analysis of mtCU components in lysates from *Quadriceps* muscle harvested from mice with whole-body depletion of Frataxin (FXN KD) (68). Fxn is a mitochondrial protein required for Fe-S cluster biogenesis. *Right*: Representative Western blots. *Left*: Quantification (mean ± s.e.m.; individual points are from different mice), n=5/genotype; *** p<0.01, unpaired t-test. **(B)** Original analysis of proteomics data from mouse heart, extracted from the mass spectroscopy dataset published by Kühl and colleagues (69). Samples were from 5 models of mice, each with a different cardiac-specific depletion (KO) of different mitochondrial proteins resulting in depletion of mtDNA. MCU and MICU1, as well as groups of functionally related mitochondrial protein from the outer and inner mitochondrial membrane (IMM, OMM) and the matrix, from nuclear- and mitochondrial-encoded DNA (nDNA, mtDNA), were expressed relative to levels in control mice. In all 5 models, MCU and MICU1 abundance was greater in KO compared to control, and the increase over the control was greater than increases in groups of other mitochondria proteins. Statistics: paired t-test, with indicated p values.

## DISCUSSION

Recent progress yielded the identity of proteins responsible for mitochondrial Ca^2+^ influx and efflux and, from that, a better understanding of these processes. However, a role for Pi to regulate the mitochondrial matrix Ca^2+^ chelation and to mitigate ΔΨm rundown during mitochondrial Ca^2+^ uptake remained less explored at the level of the relevant protein entity. The top candidate Pi transporter is the PiC, due to its high abundance and transport rate. Thus we focused on the PiC, and used a new model of PiC KD in murine SM. We also intended to mimic physiologic signaling conditions, and thus used rapid exposures of mitochondria to a modest [Ca^2+^], in contrast to an approach in which CaCl_2_ was added slowly to maximize Ca^2+^ buffering in the matrix (32). We find that PiC is the only relevant Pi transporter in SM mitochondria. Contrary to expectation, PiC is not required to support robust mitochondrial Ca^2+^ uptake which is in fact elevated in KD mitochondria, likely reflecting a higher mtCU abundance. PiC is however required to chelate matrix Ca^2+^ across a physiological range of total mitochondrial Ca^2+^. Thus this study establishes the PiC as a requirement for maintaining a bound fraction of Ca^2+^ in the matrix.

Investigating the broader impact of the mitochondrial Ca^2+^ phenotypes in KD mice and SM, we observe an exercise deficit and suppressed cytoplasmic Ca^2+^ response to tetanic stimulation, but no or minimal changes in extra-mitochondrial Ca^2+^ handling proteins and maximal SM force generation in younger KD mice. Testing for a causal link between the mitochondrial Ca^2+^ phenotype and the exercise defect in these younger mice, MCU downregulation in KD SM worsens exercise capacity. Finally, we show that elevated mtCU capacity is not unique to PiC KD but, rather, occurs in striated muscle with major mitochondrial dysfunction from a variety of causes. Thus elevated mitochondrial Ca^2+^ uptake and [Ca^2+^]m are a net benefit to PiC KD SM, and may also be so for striated SM harboring dysfunctional mitochondria more generally, at least at an early phase of this dysfunction.

### New model of PiC KD in SM

To investigate the role of the PiC in mitochondrial Ca^2+^ handling, PiC was depleted in mouse SM. As the first report of this model, we discuss here some of its broad features. Bioenergetics analyses in isolated PiC KD SM mitochondria revealed the expected biochemical defect, namely a Pi limitation on maximal oxphos. Yet oxphos of KD mitochondria was non-negligible, at ∼30% of Ctrl. There are other possible transporters of Pi across the IMM besides PiC, namely SLC25A10/23/24/25. In a proteomics analysis of SM mitochondria, SLC25A3/10/24/25 (but not SLC25A23) were detected. In the Ctrl, SLC25A3 (PiC) accounted for ∼98% of the total expression of all known possible Pi transporters (**SFig1**). In the KD, levels of SLC25A10/24/25 were unchanged (**SFig1**). Might the combined expression of SLC25A10/24/25 support Pi transport in KD mitochondria? If it would, transport rate and capacity would be low. First, the transport rate of SLC25A23/24 (70) is an order of magnitude slower than that of PiC (100-350 mmol/min/g (71)). Second, while SLC25A10’s transport rate is closer to that of PiC (∼60 mmol/min/g (72)), SLC25A10 is poorly expressed in SM (**SFig1B**). Thus, PiC is likely the major Pi transporter in SM mitochondria, and the residual PiC in KD mitochondria provides enough Pi to support oxphos that is disproportionately greater than that expected from the residual PiC protein level.

As to the impact of PiC depletion at the organ level, KD mice remain healthy into adulthood, with intact SM mass, minimal loss of maximal SM force generation, but with some exercise deficit. Similarly, mice with >90% PiC depletion in the heart are initially healthy, with normal heart size and function for several weeks after PiC loss (41). An explanation for the fairly normal organ mass and function in these models of major PiC loss can be compensations, such as higher ATP generating capacity capacity, evidenced in our study by oxphos that was only ∼30% lower than Ctrl in muscle fibers vs. 70% lower in isolated mitochondria (Fig1E vs. Fig1D), and inferred in PiC depleted heart by an initially near-normal tissue level of ATP (41). Yet the eventual outcome of major PiC loss in these models is substantial organ dysfunction: an exercise deficit with KD in SM (present study) and cardiac hypertrophy and dysfunction with PiC KD in heart (41). These phenotypes resemble the profound cardiac and skeletal myopathies described in humans with SLC25A3 variants (73–75). These organ-level observations provide useful biological contexts. The progression from modest to severe exercise phenotype in PiC KD mice provides a context to understand what can drivers or modify the progression from compensation to pathology. The relatively uncomplicated phenotype of young adult mice with SM PiC KD provides a straightforward model for studying PiC’s role in mitochondrial Ca^2+^ handling.

### Requirement for PiC to chelate matrix Ca^2+^ across a range of Ca^2+^ uptake in SM mitochondria

We evaluated [Ca^2+^]m in relation to mitochondrial Ca^2+^ uptake in the SM mitochondria, by loading mitochondria with FuraFF to measure [Ca^2+^]m, as done previously (32, 33), then simultaneously monitoring FuraFF and the buffer [Ca2+] using Rhod-FF. In KD mitochondria, we observe a greater elevation in [Ca2+]m with Ca2+ uptake, even when Ca2+ uptake in the KD was experimentally lowered to the Ctrl level (Fig2E). Elevated [Ca2+]m is also apparent when the average [Ca2+]m from each mitochondrial preparation is plotted against the corresponding Ca2+ uptake up to 50 nmol/mg mitochondrial protein (Fig2D). Thus Ca2+ chelation is less in KD mitochondria, and this is evident across a range of Ca^2+^ uptake that mitochondria can experience in vivo. We also note that, in KD mitochondria, the higher values of [Ca^2+^]m during Ca^2+^ uptake may underestimate the nmole increase in [Ca^2+^]m if Ca^2+^ uptake in KD mitochondria is accompanied by a greater increase in matrix volume (76).

Substantial lack of matrix Pi would be the straightforward explanation for the lesser Ca^2+^ chelation in KD mitochondria. However other factors could impact [Ca^2+^]m. We can rule out some of these, namely differences in baseline matrix Pi (mitochondria were pre-depleted of endogenous Pi), in Ca^2+^ efflux (CGP was included), and in mPTP opening (mPTP open probability was low, likely because ATP-Mg^2+^ was included in the incubation medium). We can also exclude pH changes. Changes in matrix pH can influence [Ca^2+^]m; alkalinization favors Pi-Ca^2+^ aggregation, thereby lowering the [Ca^2+^]m, and acidification the opposite. Though we did not measure matrix pH, a relative alkalinization resulting from Ca^2+^ uptake would be expected in KD vs. Ctrl mitochondria due to the greater depolarization in the KD during Ca^2+^ uptake (Fig3) and consequent redistribution of H^+^ from matrix to IMS/cristae space. Yet, [Ca^2+^]m is elevated in the KD. Thus any effect of alkalinization was more than offset by less availability of a species that binds Ca^2+^. On the other hand, when Ca^2+^ uptake in the KD is partially suppressed by low [RuR], matrix acidification (favoring Pi-Ca^2+^ disaggregation) might explain the lack of any decrease in [Ca^2+^]m in RuR-treated vs. untreated KD mitochondria (Fig2E). Pi-independent Ca^2+^ chelation is also possible, by cardiolipin, carboxylate anions, ATP and ADP (77–80). Yet a genotype difference in Ca^2+^ binding to ATP/ADP seems unlikely, due to the Pi pre-depletion step and ATP-Mg^2+^ in the incubation medium that would tend to equalize [ATP] between the genotypes. But we cannot exclude genotype differences in Ca^2+^ binding to cardiolipin or carboxylate anions, though we suggest that there is insufficient capacity or access to these buffers in KD mitochondria to offset the lower matrix Pi. Thus we conclude that higher [Ca^2+^]m in the KD largely reflects PiC loss and lower matrix Pi.

Though this study does not focus on the dynamics of matrix Ca^2+^ chelation (32), the relationship between mitochondrial Ca^2+^ uptake and [Ca^2+^]m in Fig2D offers insight into how PiC influences the chelation dynamics. In Ctrl mitochondria, [Ca^2+^]m varies modestly with Ca^2+^ uptake, in line with observations from Chalmers and Nicholls for mitochondrial Ca^2+^ uptake up to 120 nmol/mg mitochondria (the initial part of the trace in Figure 8A in (32)). In contrast, in KD mitochondria, there is a significant rise in [Ca^2+^]m with increasing Ca^2+^ uptake, and resembling the greater rise in [Ca^2+^]m for mitochondrial Ca^2+^ uptake from 120 to ∼480 nmol/mg mitochondria shown in (32). In (32), there was an even steeper rise in [Ca^2+^]m for Ca^2+^ uptake > 480 nmol/mg mitochondria (stated to be analogous to a Pi-free state (32)). The comparison with (32) provides a further argument that there is insufficient Ca^2+^ buffering capacity in the matrix of PiC KD mitochondria over a range of mitochondrial Ca^2+^ uptake where the matrix Ca^2+^ should be well buffered.

### Ca^2+^ uptake and mtCU abundance are elevated with PiC KD but unlikely to be unique to PiC loss

Surprisingly, Ca^2+^ uptake is greater in KD mitochondria (Fig2ABD), and appears sustainable over several 10 µM CaCl_2_ pulses (Fig3D). Mouse heart mitochondria depleted of PiC by >90% also showed robust Ca^2+^ uptake, though the magnitude of the uptake was not evaluated (41). Mitochondrial Ca^2+^ uptake has been shown to be facilitated by other anions such as ATP (33). We included ATP-Mg^2+^ in the incubation medium, and its removal increased mitochondrial Ca^2+^ uptake (compare Fig3D and E), as has also been shown (81). Our system did not contain any other candidate anion. Furthermore, any major contribution from an alternate Pi transporter seems unlikely, as discussed above. We note that, though Pi-dependence of Ca^2+^ uptake (33) was evident in Ctrl mitochondria and, to a lesser extent, in the KD, substantial Ca^2+^ uptake was evident even in the absence of Pi in the Ctrl (Fig2B).

To investigate the greater Ca^2+^ uptake in KD mitochondria, we checked the expression level of the mtCU components and the assembled complex. These are increased (Fig4), and the individual components are greater by a similar magnitude as the greater Ca^2+^ uptake (∼40%). Transcript levels are not higher (SFig7), suggesting a mechanism downstream from transcription. We surveyed the literature, tested another mitochondrial dysfunction model in our lab (loss of Frataxin), and performed an additional analysis of published proteomics data. These efforts revealed other instances where depletion of a mitochondrial protein from murine striated muscle is accompanied by greater MCU protein abundance (50, 67, 69, 82) and also MICU1 (68) (Fig6). Furthermore, in a model of Drp1 depletion in SM, MCU mRNA was measured and found to be unchanged (50). Thus elevated mtCU capacity may be a general feature of striated muscle in which mitochondrial function is substantially altered. A recent study proposed a Complex I-dependent mechanism leading to higher MCU protein (82). Complex I was not defective in PiC KD mitochondria, evidenced by higher FCCP-driven JO_2_ in KD mitochondria (Fig1D). Thus, the higher mtCU expression in PiC KD mitochondria may require further understanding. Regardless of the mechanism, an implication is that substantially altered mitochondrial function, such as PiC loss, in striated muscle can (indirectly) regulate mtCU abundance and thus the capacity for mitochondria to take up Ca^2+^.

### Suppressed cytoplasmic Ca^2+^ response to a tetanus in PiC KD muscle

Cytoplasmic Ca^2+^ signals can be shaped by mitochondrial Ca^2+^ uptake, inclusing in SM (64, 65). Given the mitochondrial Ca^2+^ phenotype in KD SM, it was of interest to investigate cytoplasmic Ca^2+^ signaling, which we did using muscle fibers stimulated with ST or RTS. The only change observed in KD fibers is a lower amplitude of the cytoplasmic Ca^2+^ response to RTS (Fig5ABFG). In younger mice, this occurs without any change in the protein level of major non-mitochondrial Ca^2+^ handling proteins, suggesting that none of these aspects of Ca^2+^ handling are responsible (Fig5C)(though post-translational modifications remain possible). The blunted response to RTS might instead reflect the greater uptake of Ca^2+^ by PiC KD mitochondria, which could lead to less Ca^2+^-mediated positive feedback on RYR1 (83). In support, muscle fibers overexpressing MCU tended to have a smaller cytoplasmic Ca^2+^ response to a KCl pulse (7). The greater suppression of the cytoplasmic Ca^2+^ response to RTS in SM from older KD mice (Fig5FG) was associated with the same increase in mtCU protein abundance as in younger but also featured a profound loss of RYR1 and Cav1.1 protein (Fig5I). So it seems reasonable to attribute the further deterioration of the cytoplasmic Ca^2+^ response to an emergant limitation in Ca^2+^ availability to the cytoplasm. Dramatically lower RYR1 abundance together a blunted cytoplasmic Ca^2+^ response to a tetanus were also shown in Drp1-depleted SM (50). To our knowledge, the present study is the first to describe a defective a cytoplasmic Ca^2+^ response without changes in the abundance of extra-mitochondrial Ca^2+^ handling proteins, though we cannot rule out other types of regulation, e.g., oxidation or nitrosylation that can functionally impact RYR1 (84, 85).

### Elevated mtCU protects exercise capacity in younger PiC KD mice

Given the Ca^2+^ phenotypes and oxphos deficit in KD SM, we checked SM function by measuring running capacity on a treadmill with a slow speed ramp-up to provide an oxphos-demanding workload. Almost all Ctrl mice completed the 90-min protocol whereas none of the KD mice did. Younger (∼13-wk-old) KD mice ran for just over half the time (Fig5D). But the older (23-25-wk-old) KD mice could barely run (Fig5E). Younger and older KD mice had a similar oxphos capacity in SM fibers, but the older KD mice showed the striking diffence from their younger KD counterparts of a marked loss, in SM, of RYR1 and Cav1.1 protein and major deterioration in the Ca^2+^ response to RTS. Drp1-depleted SM also showed a substantial lower RYR1 protein level (50). We propose that the profound cytoplasmic Ca^2+^ phenotype in older KD mice was sufficient to severely limit exercise.

Younger KD mice retained some exercise capacity, though it was less than for Ctrl mice. In the youngfer KD mice, the prominent mitochondrial Ca^2+^ phenotypes and blunted cytoplasmic Ca^2+^ response to RTS were the only (obvious) Ca^2+^ phenotypes in their SM, and maximal SM force generation and fatiguability were minimally changed from Ctrl. So we asked if the mitochondrial Ca^2+^ phenotypes might influence the exercise capacity of KD mice and, to that end, developed a model of inducible MCU depletion in PiC KD SM. Tam-treating PiC-MCU double floxed Cre+ mice lowered the MCU protein level (measured 4 wks post final Tam injection) to ∼75% of Ctrl, and thus mainly reversed the rise in MCU protein. Because MCU protein would be expected to be turned over within 4 weeks (ref), the large residual MCU protein in KD SM may reflect slower turnover. Lowering MCU in PiC KD SM caused exercise capacity to be lower and and post-run lactate to be higher compared to PIC KD mice (Fig6). Though the MCU decrease likely impacted multiple processes in KD SM, these outcomes indicate that the greater mtCU abundance in KD muscle is a net advantage, and may reflect, at least partially, a beneficial effect of elevated [Ca^2+^]m on mitochondrial metabolism. Furthermore, because several models of oxphos impairment in striated muscle show evidence for elevated mtCU abundance (Fig7), we propose that, more generally, higher mtCU expression can buffer some of the pathology in striated SM with major mitochondrial dysfunction, at least when Ca^2+^ availability to the cytoplasm is sufficient to support exercise.

## MATERIALS and METHODS

All Materials and Methods have been provided in a Supplemental File.

## Supporting information

Detailed Materials and Methods

## ACKNOWLEDGEMENTS

We thank Jeffery Molkentin, PhD (Cincinnati Children’s Hospital Medical Center) for the PiC^fl/fl^ mice, and John Elrod, PhD (Temple University) for the MCU^fl/fl^ mice. Studies were funded by Department of Defense PR161246 and NIH R01 GM123771 (to E.L.S. and G.H.), NIH R01 GM146116 (E.L.S.), NIH R01 NS132056 and GM151536 (G.H.). We thank Hsin-Yao Tang, PhD (Wistar Institute) for performing the mass spectrometry proteomics analysis.

---

**Supplemental Figure 1:**
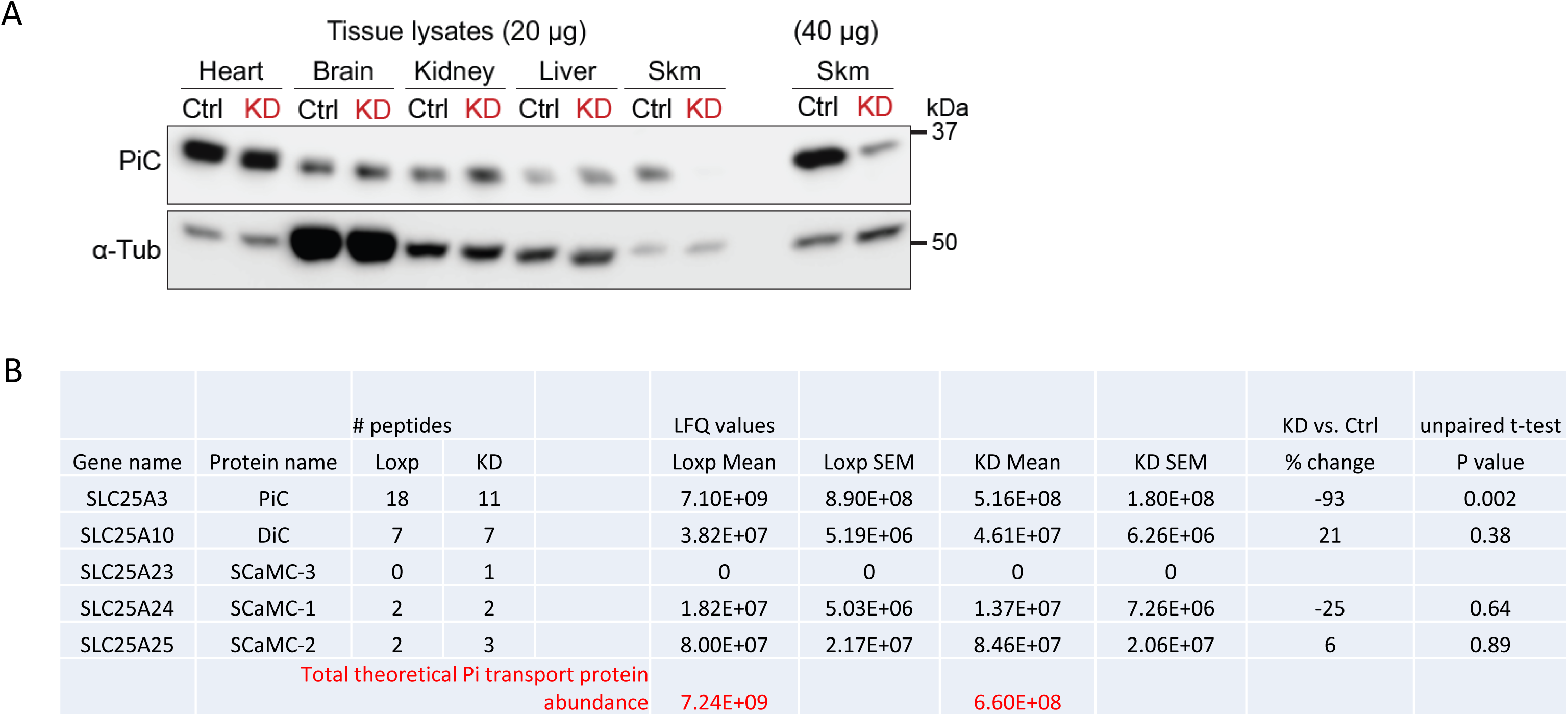
PiC depletion is confined to SM in PiC KD mice, and PiC is the dominant Pi transporter in mouse SM. (Accompanies Figure 1) **(A)** Representative Western blot of PiC in lysates prepared from different tissues harvested from 13-week old Ctrl and PiC KD mice (n=1/genotype) treated with Tam for 4 days starting at 9 weeks of age. α-Tubulin (α-Tub) was used as the loading control. For skeletal muscle (skm), *Quadriceps* was used; two different amounts of protein were loaded to reveal residual PiC protein. **(B)** Mass spectroscopy label-free based proteomics analysis of isolated SM mitochondria. Values are mean (s.e.m.; n = 3 mice/genotype) LFQ (label-free quantitation) values were normalized using the MaxLFQ algorithm (86) allowing comparison among samples. In Ctrl SM mitochondria, the PiC represents ∼96% of the theoretical Pi transport capacity.

**Supplemental Figure 2:**
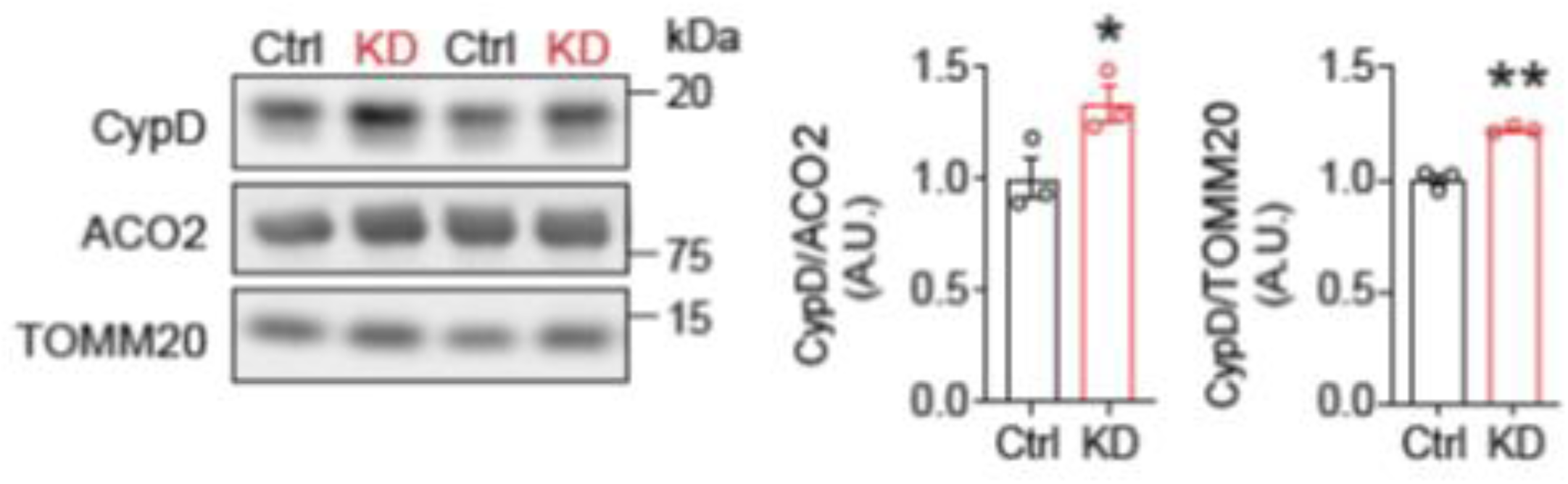
Increased CypD protein expression in PiC-deficient SM mitochondria. (Accompanies Figure 3) Representative immunoblots of CypD in SM mitochondria isolated from Ctrl and PiC KD mice. ACO2 (aconitase 2: matrix protein) and TOMM20 (outer membrane protein) were used as comparison proteins. *Right*: Averaged values relative to ACO2 and TOMM20. Bar charts: individual data points are shown, and bars represent mean ± s.e.m. Statistical comparison: unpaired t-test, \**p*<0.05, \*\**p*<0.01; n=3/genotype.

**Supplemental Figure 3:**
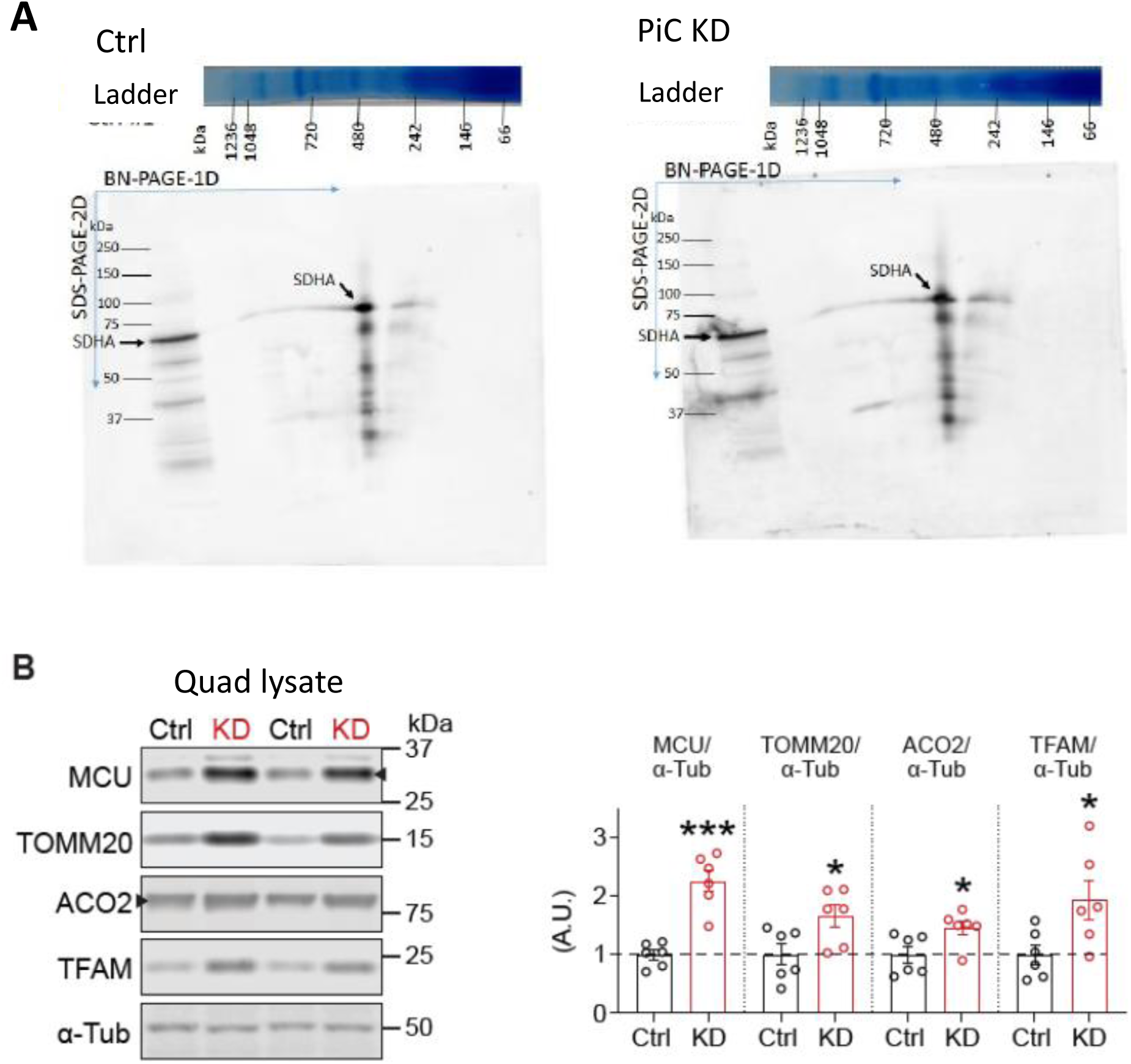
MCU is elevated relative to other mitochondrial proteins in Quadriceps lysates. (Accompanies Figure 4) (A) Nitrocellulose membranes shown in panels C, D of Figure 4 were exposed to anti-SDHA antibody, to serve as a loading control (SDHA: succinate dehydrogenase subunit A). Note that the Coomassie blue-stained gels shown in panel A also serve as loading controls. (B) Representative Western blots of MCU, TOMM20, ACO2, TFAM and α-Tubulin (α-Tub: non-mitochondrial protein and used as the loading control) in lysates from *Quadriceps* from Ctrl and PiC KD mice. *Right*: Quantification relative to α-Tubulin (n = 6/genotype).

**Supplemental Figure 4:**
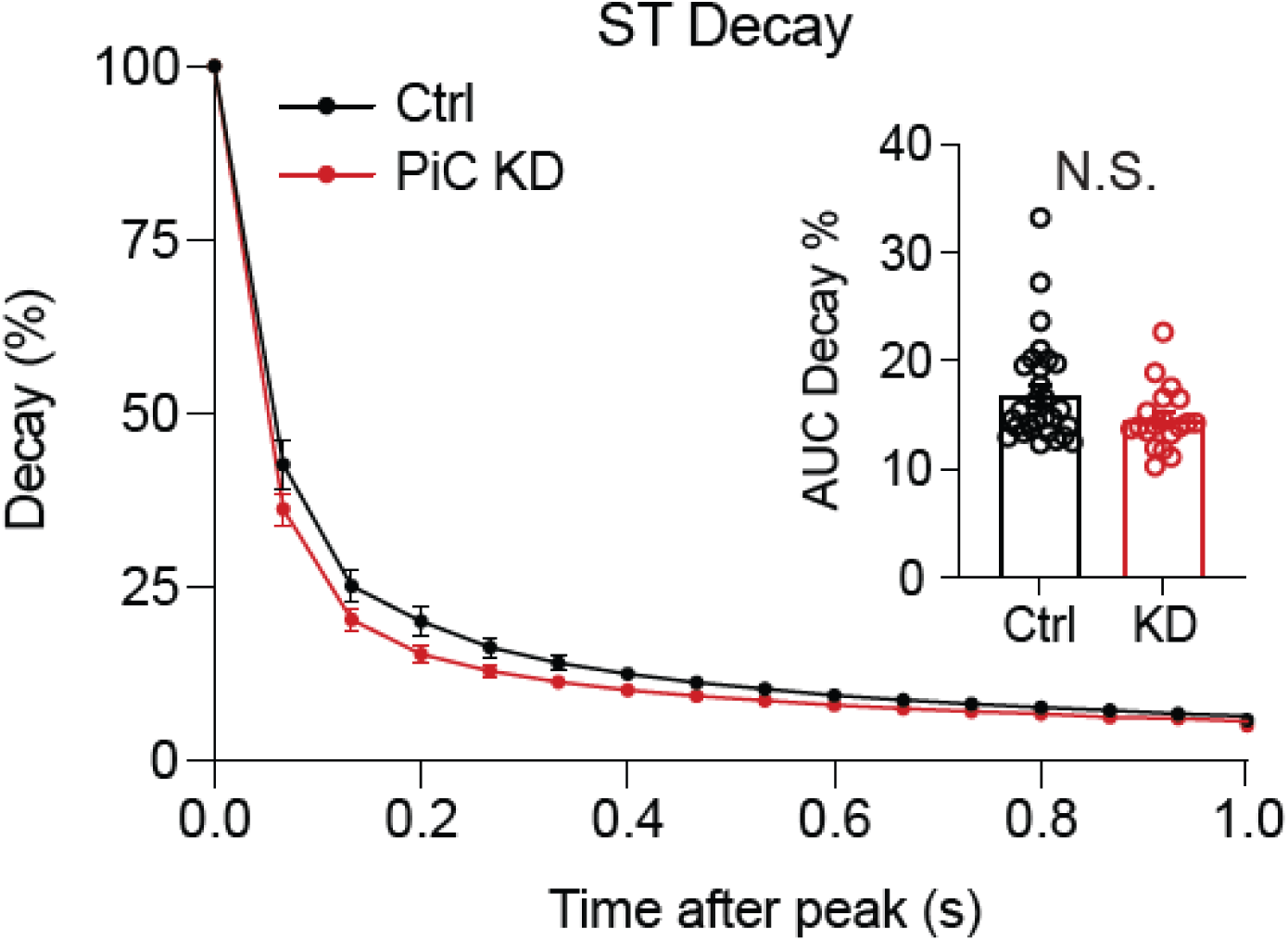
The cytoplasmic Ca2+ response shows similar decay kinetics in Ctrl and PiC KD muscle following a single twitch. (Accompanies Figure 5) Traces show average (± s.e.m.) decay of the cytoplasmic Ca^2+^ response to a single twitch delivered to *Flexor digitorum brevus* (FDB) muscle from Ctrl (black) and PiC KD (red) mice. *Inset*: Decay kinetic was quantified as area under the curve (AUC); individual data points are shown, and bars represent mean ± s.e.m. Statistical comparison: unpaired t-test, N.S.: not significant; n = 29 Ctrl and 19 PiC KD FDB.

**Supplemental Figure 5:**
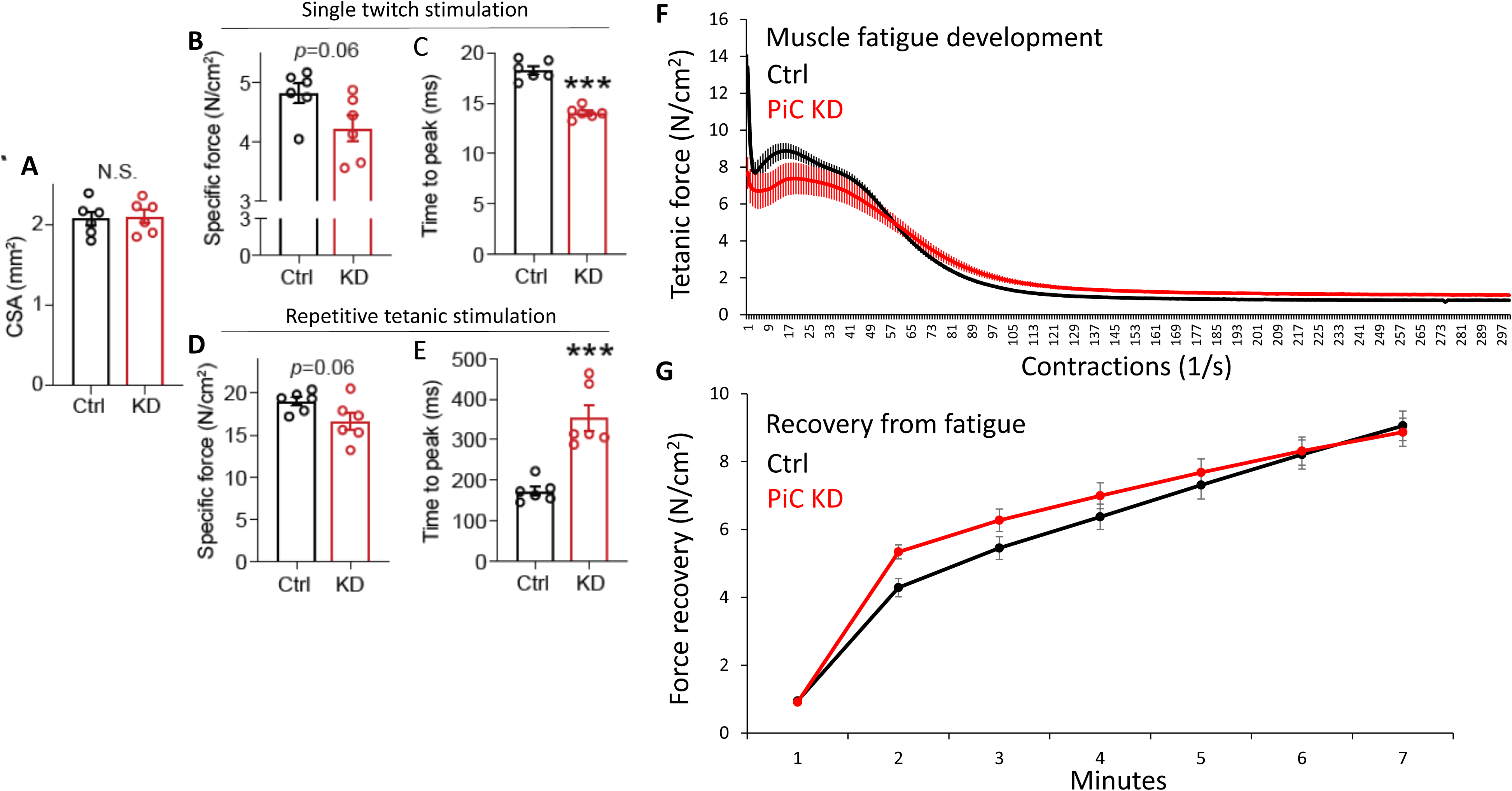
Minimal effects of PiC KD on SM force and recovery from fatigue. (Accompanies Figure 5) **(A)** Cross-sectional area (CSA) of *Extensor digitorum longus* (EDL) muscles used to evaluate force generation in Ctrl (black) and PiC KD (red) EDL. **(B) , (D)** Maximal force relative to CSA in response to single twitch stimulation (B) or repetitive tetanic stimulation (D) of EDL. **(C) , (E)** Time-to-peak during single twitch stimulation (C) or repetitive tetanic stimulation (E) of EDL. **(F)** EDL muscles were subjected to repetitive tetanic stimuli until fatigue, measured as loss of force development **(G)** Time course of force recovery after fatigue. Bar charts: individual data points are from separate mice, and bars represent mean ± s.e.m. Statistical comparison: unpaired t-test, \*\*\**p*<0.001; N.S.: not significant; n = 6/genotype.

**Supplemental Figure 6:**
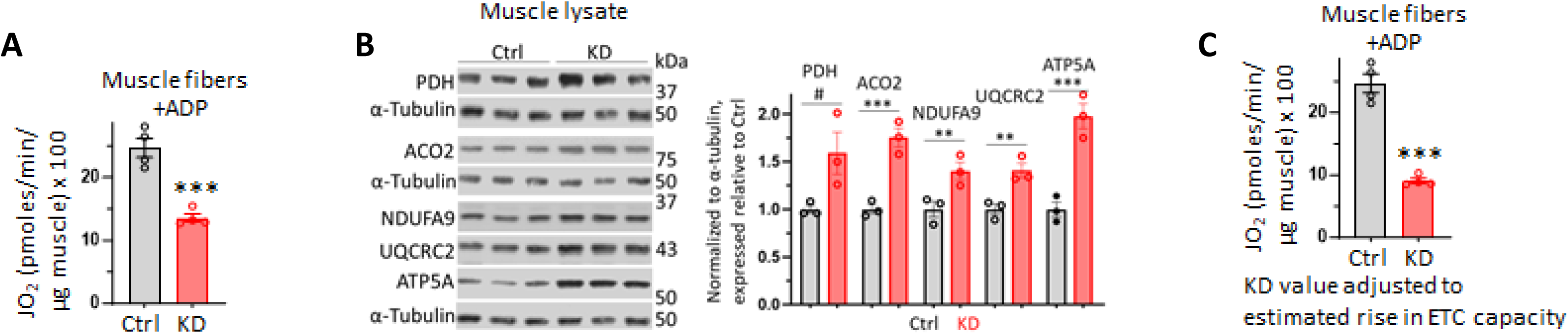
Bioenergetics status of fibers in 23-25-wk-old mice. (Accompanies Figure 5) (A) Oxygen consumption rate (JO_2_) measured in permeabilized fibers from *Extensor digitorum brevis* muscle supplied with saturating pyruvate/malate (10 mM/5 mM) and ADP (3 µM). Values were normalized to the wet weight of the muscle, measured after each series of JO_2_ experiments; n=4/genotype. (B) Western blot showing protein expression of major substrate oxidation and ETC proteins in *Quadriceps* lysate; each lane shows data from a different mouse. *Right*: Quantification, n=3/genotype. PDH: pyruvate dehydrogenase (E2 subunit). NDUFA9: Complex I subunit. UQCRC2: Complex III subunit. ATP5A: ATP synthase subunit. (C) JO_2_ values from panel E, with values from the KD adjusted for a 30% increase in substrate oxidation and ETC protein abundance, based on data from panel H. Bar charts: individual data points are shown, and bars represent mean ± s.e.m. Statistical comparison: unpaired t-test, #*p*=0.06; \*\**p*<0.01; \*\*\**p*<0.01; N.S.: not significant.

**Supplemental Figure 7:**
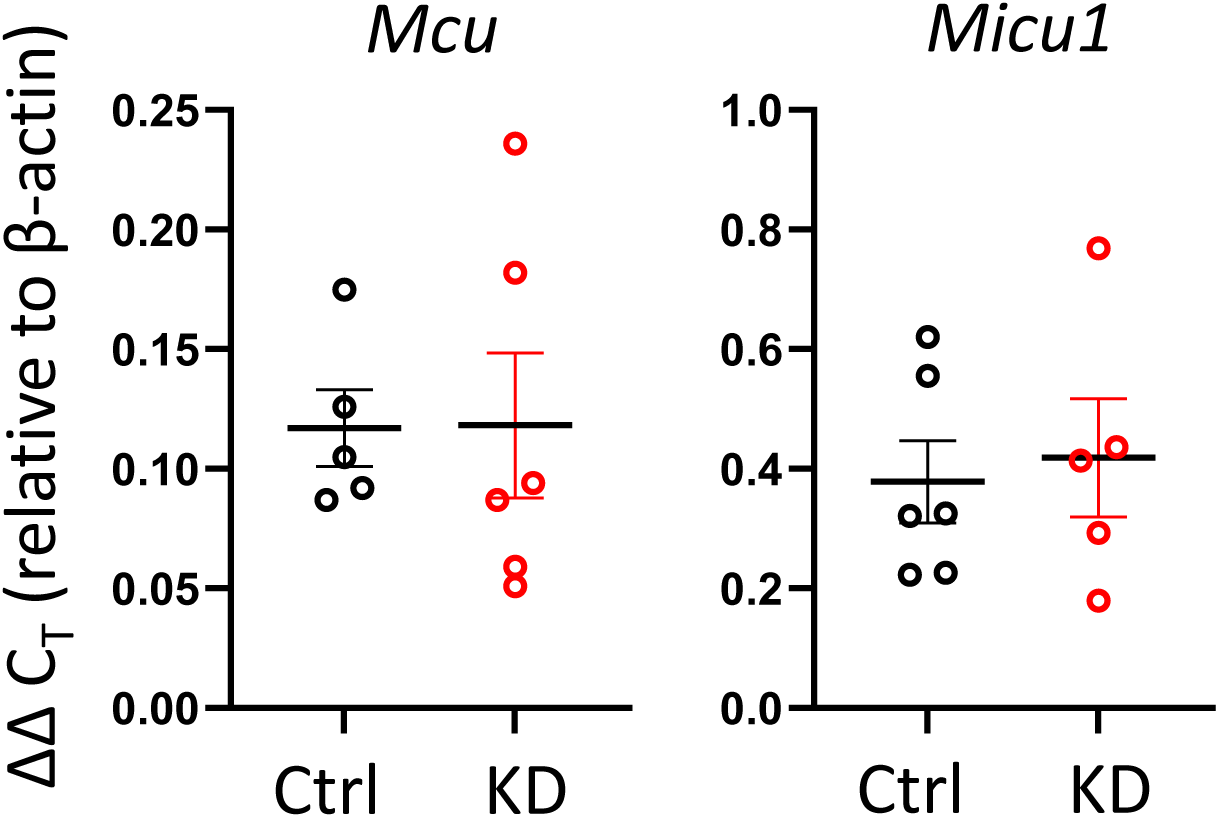
No change in Mcu or Micu1 in PiC-depleted SM. (Accompanies Figure 7) Expression of mRNA (*Mcu* and *Micu1*) in *Quadriceps* muscle. RNA was extracted from Quad muscle, from which cDNA was made, then transcript levels were measured by qPCR. Values for Mcu and Micu1 were expressed relative β-actin, using the ΔΔC_T_ method. Points and error bars represent mean ± s.e.m from n = 5-6 /genotype.

## Notes

### Competing Interest Statement

The authors have declared no competing interest.

