## Supplementary material for "Mitochondrial phosphate carrier-dependence of mitochondrial calcium chelation and respiration in skeletal muscle": Detailed Materials and Methods

**Mouse model.** Mice were used according to mandated guidelines and protocols of care and use, approved by the Thomas Jefferson University Institutional Animal Care and Use Committee (IACUC). Mice were housed at 22 °C, under a standard 12-hour light/12-hour dark cycle. Only male mice were used for experiments. Mice with specific PiC-loss in the skeletal muscle (SM) were generated by crossing mice with floxed PiC alleles (1) with mice expressing human skeletal α-actin promoter driving Cre recombinase, induced by tamoxifen (Tam) (HSA-MCM (2)). Both models were on the C57BL/6 background. Control ($PiC^{fl/fl}$) and PiC KD ($PiC^{fl/fl}$ + Cre) mice were injected intraperitoneally with Tam (25 mg/kg/mouse, i.p.) for 4 consecutive days starting when mice were 9 weeks old. Mice were sacrificed 3 and 13-15 weeks after the final Tam injection. We however noticed that HSA-MCM-$PiC^{fl/fl}$ mice that were not treated with Tam showed substantial depletion of PiC in SM. Re-deriving the colony using new founders yielded the same whereas HSA-MCM mice from our colony that were crossed with other floxed models showed the expected Tam-dependence (not shown). Thus, the Tam-independent PiC loss was likely a peculiarity of the HSA-MCM-$PiC^{fl/fl}$ model. We determined that 4 days of Tam treatment further decreased PiC protein in the soleus muscle. Thus we systematically treated 9-week-old HSA-MCM-$PiC^{fl/fl}$ mice and control $PiC^{fl/fl}$ mice with Tam, and will refer to them as PiC KD and Ctrl mice, respectively.

SM-specific depletion of both PiC and MCU was achieved by generating mice harboring both $PiC^{fl/fl}$ and $MCU^{fl/fl}$ and 1 copy of HSA-MCM. $MCU^{fl/fl}$ mice used were from (3). We determined that MCUfl/fl was indeed under control of the HSA-MCM promoter (i.e., tamoxifen was required for MCU depletion). $PiC^{fl/fl}$-$MCU^{fl/fl}$-HSA-MCM mice and $PiC^{fl/fl}$-$MCU^{fl/fl}$ mice were treated with Tam as described above.

**Isolation of SM mitochondria.** SM mitochondria were isolated as described in (4). All steps were performed at 4°C or on ice. SM from mouse forelimb and hindlimb was dissected and placed in basic medium (BM) containing 140 mM KCl, 20 mM HEPES, 5 mM $MgCl_2$, 1 mM EGTA, pH 7.0. Muscle was cleaned of fat and connective tissue, minced and placed in 30 mL of homogenization medium (HM) composed of BM with the following additions: 1 mM $Mg^{2+}$-ATP, 2 mM EGTA and 1% fatty acid free BSA (w/v) containing two units of protease from *Bacillus licheniformis* (Sigma, P5380) per muscle weight (g). The tissue was homogenized using a glass Potter-Elvehjem homogenizer with a Teflon pestle (15 passes at 500 r.p.m.). The tissue suspension was centrifuged at 500 x g for 10 min, then supernatant was collected and centrifuged at 9,500 x g for 8 min. The resulting pellet was resuspended and incubated in BM, on ice, for 5 min. Then, samples were centrifuged at 500 x g for 10 min, then the supernatant was filtered through a 70 μM cell strainer (Falcon, 352250) then spun at 9,500 x g. The final pellet was resuspended in BM (to have protein concentration ~25 mg/ml). Protein concentration was determined by BCA assay (ThermoFisher Scientific, 23225).

**Fluorometric measurements in suspensions of isolated mitochondria.** For fluorometric measurements, SM mitochondria were depleted of endogenous phosphate. To achieve this, a sample of mitochondria was pelleted then resuspended in BM with the following additions: 0.75U/mL hexokinase (Roche, 11426362001), 1 mM glucose, 0.5 mM ADP, 1 mM $MgCl_2$, 5 mM malate and 5 mM pyruvate, and then incubated for 10 min at 35 °C. After that, the sample was centrifuged at 9,000 *g* for 10 min, and the resulting pellet was resuspended in BM. For fluorometric measurements of mitochondrial matrix $Ca^{2+}$ concentration ($[Ca^{2+}]_m$), SM mitochondria were loaded with $Ca^{2+}$ indicator dye by incubation with 20 µM Fura-FF free acid, AM (AAT Bioquest, 21027) and 0.3 % pluronic acid (Invitrogen, P6867) at 35 °C for 30 min. After that, the suspension was centrifuged at 9,000 *g* for 10 min at 4 °C and the resulting pellet was resuspended in BM. The extramitochondrial $Ca^{2+}$ concentration ($[Ca^{2+}]_c$), was assessed using 500 nM Rhod-FF, tripotassium salt (AAT Bioquest, 21076) when measured simultaneously with the $[Ca^{2+}]$m. $[Ca^{2+}]_c$ was monitored using 1µM Fura2-FF pentapotassium salt (AAT Bioquest, 21025) when measured simultaneously with the $\Delta\Psi_m$ with 1.5µM TMRM (Invitrogen, T668).

All fluorometric measurements were performed using a running buffer (RB) composed of an intracellular medium (ICM) containing 120 mM KCl, 10 mM NaCl, 1 mM $KH_2PO_4$, 20 mM Tris-HEPES, pH 7.2, supplemented with protease inhibitors (1 µg/mL each of leupeptin, pepstatin and antipain), 2 µM thapsigargin, 2 mM Mg-ATP (unless otherwise stated), 0.5 µL/mL of 1 M KOH and 6 mM of each pyruvate and malate. For each run, 375 µg of mitochondria was used. CGP37157 (20 µM) was also included, to inhibit $Na^+$-dependent $Ca^{2+}$ efflux. Fluorescence was monitored in a fluorimeter (Delta-RAM, Photon Technology International) using 340 nm, 380 nm excitation and 500 nm emission for fura2FF and 540 nm excitation and 580 nm emission for Rhod-FF/TMRM. After the addition of $CaCl_2$ pulses, 6.7 µM FCCP was added to dissipate the $\Delta\Psi_m$, followed by an addition of 4 µM ionomycin. Calibration of the $[Ca^{2+}]$ signal was carried out at the end of each measurement, by adding 1 mM $CaCl_2$ followed by 10 mM EGTA/Tris, pH 8.5.

**Bioenergetics analyses in isolated skeletal mitochondria**. $O_2$ consumption ($JO_2$) was measured using the Seahorse XFe24 Analyzer (Agilent Technologies, CA, USA). Isolated mitochondria were studied as described (5, 6). Each well of the custom microplate contained 5 µg of mitochondria suspended in mitochondria assay medium (MAS): 70 mM sucrose, 22 mM mannitol, 10 mM $KH_2PO_4$, 5 mM $MgCl_2$, 2 mM HEPES, 1 mM EGTA, 0.2% fatty acid free BSA, pH 7.4 at 37°C. The microplate was centrifuged at

2,000 *g* for 20 min at 4°C. The µg of mitochondria used was optimized to obtain a linear $O_2$ *vs*. time in all conditions. Mitochondria were energized using malate-pyruvate (5 mM/10 mM). Maximal oxidative phosphorylation was measured using saturating [ADP] (2.8 mM). Maximal leak $JO_2$ was measured in the presence of the ATP synthase inhibitor oligomycin (2.5 µg/mL). Maximal uncoupled $JO_2$ was measured using the chemical uncoupler FCCP (1 µM).

**Immunoblot analysis**. For western blot analysis, SM was rapidly frozen in liquid nitrogen. Tissue was homogenized on ice in a lysis buffer containing: 150 mM NaCl, 25 mM HEPES, 2.5 mM EGTA, 1% Triton 100X, 1% Igepal (10%), 0.10% SDS, 0.1% Deoxycholate, 10% Glycerol, protease inhibitor (Roche 11873580001) and phosphatase inhibitor cocktail (200 mM sodium fluoride, 200 mM, imidazole, 115 mM sodium molybdate, 200 mM sodium orthovanadate, 400 mM sodium tartrate dihydrate, 100 mM sodium pyrophosphate and 100 mM β-glycerophosphate), using a glass/Teflon homogenizer at 500 r.p.m. Muscle lysates were incubated at 4°C for 45 min, then centrifuged at 18,000 *g* for 20 min. Protein concentration was measured from the supernatant, by BCA assay. For western blot analysis from isolated mitochondria, lysis was obtained using RIPA buffer (Sigma-Aldrich R0278) containing protease and phosphatase inhibitors. Samples were incubated on ice for 30 min and vortexed every 5-10 min, then lysates were centrifugated at 18,000 x g for 10 min. From the supernatant, protein concentration was measured by BCA assay. Primary antibodies were used for overnight incubation diluted in TBS-T 1% (20 mM Tris, 0.9 % NaCl, 0.1 % Tween 20, pH 7.4) and are listed below.

Primary Antibodies used in immunoblot analysis

| Antibody | Catalog Number | Dilution | Host |
|---|---|---|---|
| SLC25A3/PiC | Custom made | 1:1000 | Rabbit |
| α-Tubulin | CST-2215S | 1:1000 | Rabbit |
| Cyclophilin D | Abcam (Ab)-110334 | 1:10000 | Mouse |
| MCU | HPA016480 (Sigma-Aldrich) | 1:1000 | Rabbit |
| TOM20 | ProteinTech 11802 | 1:1000 | Rabbit |
| Aconitase 2 | Ab-129069 | 1:1000 | Rabbit |
| mtTFA/TFAM | Santa Cruz (SC)-23588 | 1:500 | Mouse |
| EMRE | Bethyl A300BL19208 | 1:1000 | Rabbit |
| MICU1 | HPA037479 | 1:500 | Rabbit |
| SERCA1 | Custom made, YenZym | 1:1000 | Rabbit |
| Calsequestrin | Ab-3516 | 1:1000 | Rabbit |
| Calreticulin | Cell Signaling Technology (CST)-2891 | 1:1000 | Rabbit |
| RYR1 | Developmental Studies Hybridoma Bank (DHSB)-34C | 1:100 | Mouse |
| Cav1.1 | Thermo-Fisher MA3-920 | 1:1000 | Mouse |
| GRB2 | SC-8034 | 1:1000 | Rabbit |
| Vinculin | CST-13901 | 1:1000 | Rabbit |

**Blue native (BN) PAGE and second dimension electrophoresis.** BN-PAGE was conducted essentially as in (7). After BCA, 100 µg of isolated SM mitochondria were extracted with 4% digitonin (D141-100MG,Sigma) in extraction buffer ( 30mM HEPES, 12% glycerol, 150mM potassium acetate, 2mM aminocaproic acid, 2mM EDTA disodium salt, protease inhibitor tablet, pH 7.4). To the supernatant was added 1ul of 1:400 diluted G-250: extraction buffer and the samples were loaded on a NativePAGE™ NovexR 3–12% Bis-Tris gel. The gel was run overnight (on ice, 30V) in 1x NativePAGE™ Running buffer (as anode buffer) and 1x dark blue cathode buffer for 1h then switched to 1x light blue cathode buffer for the rest of the run. Each sample was run in duplicate; one of the lanes was Coomassie blue-stained and the second lane was run on a second dimension SDS gel. The lane for the second dimension was excised, then equilibrated in an Equilibration (Eq) buffer containing 50 mM Tris, 6 M Urea, 30% glycerol, 2% SDS, pH 8.5, for 5 min, then switched to Eq. buffer + 200mM DTT for 30 min, followed by Eq. buffer + 135mM iodoacetamide for 15 min and 2x Eq. buffer for 15min incubation. The lane was then inserted into NuPAGE™ 4-12% Bis-Tris Gel with 2D well and run in 1x MOPS running buffer. Protein was transferred onto a 0.2 µm nitrocellulose membrane, which was used for protein detection with antibodies.

**Proteomics.** Label-free proteomics on samples of isolated mitochondria were performed as in (5). Samples (25 µg) of isolated SM mitochondria were run into a NuPAGE 10% Bis-Tris gel for a short distance. The entire gel lane was excised and digested with trypsin. Liquid chromatography tandem mass spectrometry (LC-MS/MS) analysis was performed using a Q Exactive HF mass spectrometer (Thermo Scientific) coupled with an UltiMate 3000 nano UPLC system (Thermo Scientific). Samples were injected onto a PepMap100 trap column (0.3 x 5 mm packed with 5 µm C18 resin; Thermo Scientific). Peptides were separated by reversed phase HPLC on BEH C18 nanocapillary analytical column (75 µm i.d. x 25 cm, 1.7 µm particle size; Waters) using a 4-h gradient formed by solvent A (0.1% formic acid in water) and solvent B (0.1% formic acid in acetonitrile). Eluted peptides were analyzed by the MS set to repetitively scan m/z from 400 to 1800 in positive ion mode. The full MS scan was collected at 60,000 resolution followed by data-dependent MS/MS scans at 15,000 resolution on the 20 most abundant ions exceeding a minimum threshold of 20,000. Peptide match was set as preferred, exclude isotope option and charge-state screening were enabled to reject unassigned and single charged ions.

Peptide sequences were identified using MaxQuant 1.6.17.0 (8). MS/MS spectra were searched against a UniProt mouse protein database (October 2020) and a common contaminants database using full tryptic specificity with up to two missed cleavages, static carboxamidomethylation of Cys, and variable Met oxidation, protein N-terminal acetylation and Asn deamidation. “Match between runs” feature was used to help transfer identifications across experiments to minimize missing values. For analysis, values were normalized using the MaxLFQ algorithm (8) to allow comparison among samples.

**Quantitative polymerase chain reaction.** Total RNA was extracted from mouse *Quadriceps* according to the manufacturer's instructions using Animal Tissue RNA Purification Kit (Norgen Biotek Corp, 25700). RNA concentration was measured by Nanodrop One (ThermoFisher Scientific). RNA was reverse transcribed using oligo(dT) primers and SuperScript III (Invitrogen, 18080-051). qPCR reactions were performed using SYBR™ (Invitrogen, 4472908), with 20 ng cDNA/reactions, using a QuantStudio™ 5 Real-Time PCR Instrument (Applied Biosystems). Levels of mRNA were calculated relative to *Actb* (β-actin) using the ΔΔCt method. Primer sequences are listed below and have been validated against knockout controls (10).

| Primer | Forward | Reverse |
|---|---|---|
| *Mcu* | *AAAGGAGCCAAAAAGTCACG* | *AACGGCGTGAGTTACAAACA* |
| *Micu1* | *AACAGCAAGAAGCCTGACAC* | *CTCATTGGGCGTTATGGAG* |
| *Actin, beta* | *CAACACCCCAGCCATG* | *GTCACGCACGATTTCCC* |

**FDB fiber isolation.** Mouse *Flexus digitorum brevis* (FDB) isolation was performed essentially as described in (11, 12). Briefly, FDB muscles from each footpad were quickly dissected and digested with 3 mg/mL of collagenase type 2 (Worthington Biochemical Corporation, LS004194) at 37 °C for 1-2 hrs. under agitation. The collagenase solution was prepared in DMEM/F12 medium (Lonza, BE04-687F/U1) without serum. After incubation with collagenase, FDB fibers were incubated with DMEM/F12 + 10% horse serum (Gibco, 16050130) containing 2 mM glutamine, 100 U/ml penicillin and 100mg/ml streptomycin (ThermoFisher Scientific 10378016). Mechanical disaggregation of the muscle was carefully done using glass pipettes, until individual fibers were obtained, which were plated on glass bottom microwell dishes (MatTek Corporation, 801002) that had been pre-treated with Cell-Tak (Corning, 354240). Fibers were allowed to attach to the dish in an incubator at 37°C with 5% $CO_2$-humidified air for 2-3 hrs prior to cell imaging.

**$Ca^{2+}$ imaging in resting and stimulated FDB muscle fibers.** To enable electrical stimulation, FDB fibers were mounted in a perfusion chamber (Warner Instruments, RC-37FS, 64-0366) holding two parallel platinum electrodes positioned 6 mm apart. The chamber was connected to a pulse stimulator (Aurora Scientific Inc, 701C) and an oscilloscope (Tektronix, TDS 220). Recordings were performed in an extracellular medium (ECM): 0.25% BSA-extracellular medium (ECM) containing: 121 mM NaCl, 5 mM

$NaHCO_3$, 4.7 mM KCl, 1.2 mM $KH_2PO_4$, 1.2 mM $MgSO_4$, 2 mM $CaCl_2$, 10 mM glucose and 10 mM Na-HEPES, pH 7.4, at 35 °C.

To image $[Ca^{2+}]c$, fibers were loaded with 2 µM Fura-2, AM (Invitrogen, F-1221) at room temperature in 2% BSA-ECM in presence of 0.003% pluronic acid (Invitrogen, P6867) and 100 µM sulfinpyrazone (Sigma, 9509), for 20 min. After calcium dye loading, fibers were carefully rinsed with ECM, and maintained in ECM containing 75 µM BTS (Tocris, 1870) and 100 µM sulfinpyrazone at 35 °C. Images were acquired using an ImagEM EM-CCD camera (Hamamatsu) fitted to an Olympus IX81 microscope with LED source (Lambda TLED+, Sutter Instruments). Fura-2 fluorescence was detected by excitation at 340/380 nm (emission 540/50 nm). Calibration of the Fura2-AM signal was carried out by adding 1 mM $CaCl_2$, then 10 mM EGTA/Tris, pH 8.5.

FDB fibers were electrically stimulated by means of field stimulation as described (11). The repetitive tetanic stimulation (RTS) protocol was: 100 tetani, 100 Hz, for 500 ms every 2.5 s, pulse length 2 ms, at 2 V. For single twitch (ST), fibers were stimulated at 1 Hz every 60 s, pulse length 2 ms at 2V. The RTS and ST frequency was controlled by an Optogenetics TTL pulse generator (Doric, OPTG4).

**Measurement of $O_2$ consumption ($JO_2$) in permeabilized FDB muscle fibers.** Permeabilized muscle bundles were prepared and $JO_2$ measurements were performed essentially as in (12). A small amount (~25 mg) of *Extensor digitorum longus* (EDL) muscle was dissected, trimmed of connective tissue and fat, then placed in ice cold Buffer IB (Isolation Buffer: 7.23 mM $K_2EGTA$, 2.77 mM $CaK_2EGTA$, 20 mM imidazole, 20 mM taurine, 5.77 mM $MgCl_2$, 50 mM K-MES, 0.5 mM dithiothreitol; 15 mM $Na_2$Phosphocreatine pH 7.3 at 4°C). To prepare fiber bundles, the muscle was gently separated longitudinally using fine forceps. Bundles were permeabilized by a 30-min incubation in saponin (50 ug/ml) at 4°C, in Buffer RB (Respiration Buffer), then washed 3x, 5 mins each, on ice, in Buffer RB: 110 mM sucrose, 100 mM mannitol, 30 mM KCl, 60 mM K-MES, 1.5 mM EGTA, 3 mM $MgCl_2$, 10 mM $K_2HPO_4$, 20 mM HEPES, 20 mM taurine, pH 7.3 at 4°C, supplemented with 1 mg/ml BSA. Muscle bundles were maintained in Buffer RB, on ice, until use.

$JO_2$ was measured in muscle bundles (~1 – 2.5 dry weight) using an Oroboros Oxygraph-2k (Oroboros Instruments), under continuous stirring (750 rev/min), at 37°C, in 2 ml of Buffer Z. Pyruvate and malate (10 mM and 5 mM) were used as substrates, and ADP (2 mM) was used to stimulate oxphos. After each run, bundles were retrieved, carefully rinsed with $ddH_2O$, then dried overnight in a 37°C oven. $JO_2$ values were expressed relative to dry muscle weight.

***Ex vivo* physiology in EDL.** Evaluation of *ex vivo* EDL force was performed as described (13), using an Aurora Mouse 1200A System equipped with Dynamic Muscle Control v.5.415 software. Briefly, EDL muscles were dissected and maintained in constantly oxygenated Ringer's solution (100 mM NaCl, 4.7 mM KCl, 3.4 mM $CaCl_2$, 1.2 mM $KH_2PO4$, 1.2 mM MgSO4, 25 mM HEPES, 5.5 mM D-glucose) at 24 °C. The twitch stimulation protocol applied was a single 30-V stimulus with a duration of 0.2 ms. For measuring tetanic maximal force generation, the 30-V stimulus lasting 0.2 ms was repeated at a frequency of 120 Hz for 500 ms. Five minutes were allowed between two tetanic contractions to ensure muscle recovery.

**Treadmill running**. Treadmill running was performed using an Exer3/6 Treadmill (Columbus Instruments). Mice were acclimated to the treadmill over 3 consecutive days. On days 1 and 2, mice were were placed on a stationary treadmill for 10 minutes, with the shocks off. On acclimation day 3, mice ran at 5 m/min for 3 minutes, then at 10 m/min for 8 minutes, with shocks on. The test run, a slow incremental speed endurance-type run, and performed after 1 or 2 days of rest, was as follows: speed was increased from 0 to 5 m/min then increased in 2.5 m/min increments until a maximum speed of 25 m/min. Duration at the 5 – 10 m/min speeds was 12 minutes, at the 12.5 – 17.5 m/min speeds was 10 minutes, at the 20 and 22.5 m/min speeds was 6 minutes, then at 25 m/min was 4 minutes.

**Quantification and statistical analysis.** Data analysis was performed using GraphPad Prism 8.2.1 Software. Paired Student's *t* test for comparison of two means and one or two-way analysis of variance followed by Tukey post-hoc comparisons, for multiple comparisons, were performed as appropriate. P value $< 0.05$ was considered significant. For details are provided in Results and the figure legends.
